# From Closed-Set Annotation to In-Context Discovery: Biologically Grounded LLMs for Cellular Profiling

**DOI:** 10.64898/2026.10.07.757352

**Authors:** Maxence Gélard, Hakim Benkirane, Thomas Pierrot, Paul-Henry Cournède, Bernardo P. de Almeida

**Author notes:** These authors jointly supervised this work (Paul-Henry Cournède, Bernardo P. de Almeida).

## Abstract

Tumour heterogeneity reflects both cellular *identity* (lineage) and *state* (active transcriptional programs), yet computational methods generally analyse these axes separately. We introduce CellScribe, a generative framework that conditions a pretrained language model on frozen single-cell foundation model embeddings through AdaLN adapters, enabling one shared model to generate an ontology-grounded cell type or a quantified meta-program profile. Supervised multitask fine-tuning produces fine-grained typing across hundreds of cell types and outperforms generative and retrieval baselines on meta-program identification under study-holdout evaluation. We further train the model with reinforcement learning, scoring its generations against cell ontology distance and meta-program agreement, so that they align with biological knowledge rather than token agreement. To identify new cell types not present in the supervised training data, we post-train the model to read several labelled example cells in a single context, each conditioning its own text span, and infer the type of a query cell through in-context learning. CellScribe thus unifies cell identity and state in one generative interface, grounds both in biological structure, and identifies cell types outside the supervised vocabulary.

## 1 Introduction

Single-cell RNA sequencing (scRNA-seq) [Tang et al., 2009] helps resolve tumour composition and functional heterogeneity that bulk measurements average away. Its interpretation requires two complementary answers: **identity**, the cell’s lineage (e.g., a CD8 T cell), and **state**, the dynamic processes it exhibits (e.g., proliferation, hypoxia, or stress). Cells of the same lineage, even within one tumour, can differ in invasive potential, treatment susceptibility, and microenvironmental interactions. Nevertheless, identity and state are typically analysed with separate tools, and are rarely reported together.

### Defining the two tasks

*Cell type* is the lineage a cell belongs to, drawn from a controlled vocabulary: the cell ontology [Diehl et al., 2016], a hierarchy running from broad classes to fine subtypes. Identity is single-label and hierarchical, so a coarser prediction is partially correct. For *cell state*, we follow the definition introduced with the Curated Cancer Cell Atlas (3CA) by Gavish et al. [2023]. Integrating 1,163 tumour samples across 24 cancer types, they identified 41 recurrent meta-programs: patterns of coordinated gene expressions across malignant cells within tumours. These programs capture generic processes, including cell cycle, hypoxia, stress, and interferon responses, as well as epithelial-mesenchymal transition. They are grouped into 11 broader hallmarks of transcriptional intratumour heterogeneity. We therefore represent state at two resolutions: a set of active meta-programs with continuous activity scores and the dominant hallmark summarising them. Unlike cell identity, the fine-grained state is multi-label: one cell may be cycling, hypoxic, and stressed simultaneously. Per-cell activity profiles can expose co-active and rare states that tumour-level averages obscure [Barkley et al., 2022, Tirosh and Suvà, 2024].

### Limitations of current tools

Cell-type annotation still requires experts to cluster cells, inspect markers, and validate tissue-specific labels. Cross-tissue supervised models [Fischer et al., 2024] accelerate and standardise this process, but fixed output heads remain confined to predefined types. State inference is complicated by overlapping programs: ssGSEA [Barbie et al., 2009] and AUCell [Aibar et al., 2017] score signatures largely independently and are sensitive to shared genes and sparse measurements. Conversely, the NMF procedure [Lee and Seung, 1999] used to discover meta-programs fits cohorts retrospectively rather than predicting the state of a new cell. A portable model should therefore resolve overlapping programs while representing identity and state through one cross-study interface.

### Our approach

We develop CellScribe, a large language model (LLM) conditioned on single-cell embeddings that provides a common interface for identity and state. Supervised fine-tuning (SFT) teaches the model to answer both prompts with structured text. Group Relative Policy Optimisation (GRPO) then aligns these generations with cell ontology distance [Diehl et al., 2016] and meta-program agreement. Span-conditioned in-context learning (ICL) further allows the model to compare a query with labelled cells in the same prompt, a capability already widely used for text-only models [Brown et al., 2020].

### Contributions

To our knowledge, no previous model combines fine-grained cell typing and quantitative cell-state prediction within one generative interface while also supporting ontology-aware reward alignment and multi-cell prompting. We make four contributions:

- **CellScribe**, a generative single-cell model that conditions a pretrained LLM on frozen scFoundation embeddings through AdaLN-Zero modulation [Peebles and Xie, 2023], emitting anontology-grounded cell type or a quantified meta-program profile from one shared model under task-specific prompts (§3, §4).
- **Reinforcement-learning alignment** of an scRNA-seq-conditioned LLM for both cell typing and cell state. GRPO scores structured generations against cell ontology distance and meta-program agreement, improving hierarchical typing and state prediction over supervised fine-tuning (§5).
- **In-context learning with span-conditioned AdaLN**, which lets each labelled support cell modulate its own text span, so the decoder can read multiple cells in one prompt and identify cell types excluded from supervised training (§6).
- **A study-holdout evaluation on CELLxGENE and 3CA** for cell typing and cell state (§3, §4).

## 2 Related Work

### Annotators and foundation models

Seurat label transfer [Stuart et al., 2019] and CellTypist [Domínguez Conde et al., 2022] transfer reference labels through nearest-neighbour or logistic models. At scale, scTab [Fischer et al., 2024] trains a cross-tissue classifier on 22M cells and 164 types; hierarchical cross-entropy further improves out-of-distribution accuracy by incorporating the cell ontology [Cultrera di Montesano et al., 2026]. Self-supervised models such as scGPT [Cui et al., 2024], Geneformer [Theodoris et al., 2023] and scFoundation [Hao et al., 2024] instead learn reusable representations from millions of cells. These methods remain closed-set at inference and do not quantify recurrent cell-state programs.

### Ontology-aware and language-based methods

OnClass [Wang et al., 2021] jointly embeds cells and ontology terms so unseen types can be annotated by ontology proximity, while SCimilarity [Heimberg et al., 2025] learns a cross-study retrieval space; both match in embedding space rather than generate. CellWhisperer [Schaefer et al., 2025] aligns expression and text with a CLIP-style objective [Radford et al., 2021], which scores a cell against a caption but does not decode a structured type or program profile. Cell2Sentence [Levine et al., 2024] serialises ranked genes into free text, and InstructCell [Fang et al., 2025] instruction-tunes an LLM across single-cell tasks, yet neither predicts quantitative recurrent state, aligns generations with ontology or program structure, nor conditions on labelled cells.

### Cancer cell state

We adopt the 41 meta-programs and 11 hallmarks of Gavish et al. [2023] as the operational definition of cell state. These programs are recurrent: the same transcriptional states appear across tumours and cancer types [Barkley et al., 2022, Tirosh and Suvà, 2024], which is why a shared vocabulary can be predicted on new studies.

## 3 The CellScribe Model

### 3.1 Architecture

#### Overview

Let 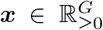 denote the expression profile of one cell over *G*=19264 genes. A pretrained single-cell RNA-seq encoder *E* maps this profile to a 3072-dimensional cell-level representation by concatenating four pooled views of its final hidden states. A learned linear layer projects this representation into the LLM hidden space, producing the conditioning embedding 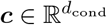 with *d*_cond_=896. In parallel, a task-specific prompt *P*_*t*_, with *t* ∈{type, state}, is tokenised and passed to a causal large language model (LLM) *π*_*θ*_. Conditioned on ***c***, the LLM generates the answer *A* = (*a*_1_, …, *a*_*T*_ ) according to *π*_*θ*_(*A*|*P*_*t*_, ***c***). We instantiate the LLM as `Qwen2.5-0.5B-Instruct` [Qwen Team, 2024] (*L*=24, *d*_model_=896) and use the 100M-parameter scFoundation encoder [Hao et al., 2024], pretrained on 50M cells (Figure 1).

**Figure 1:**
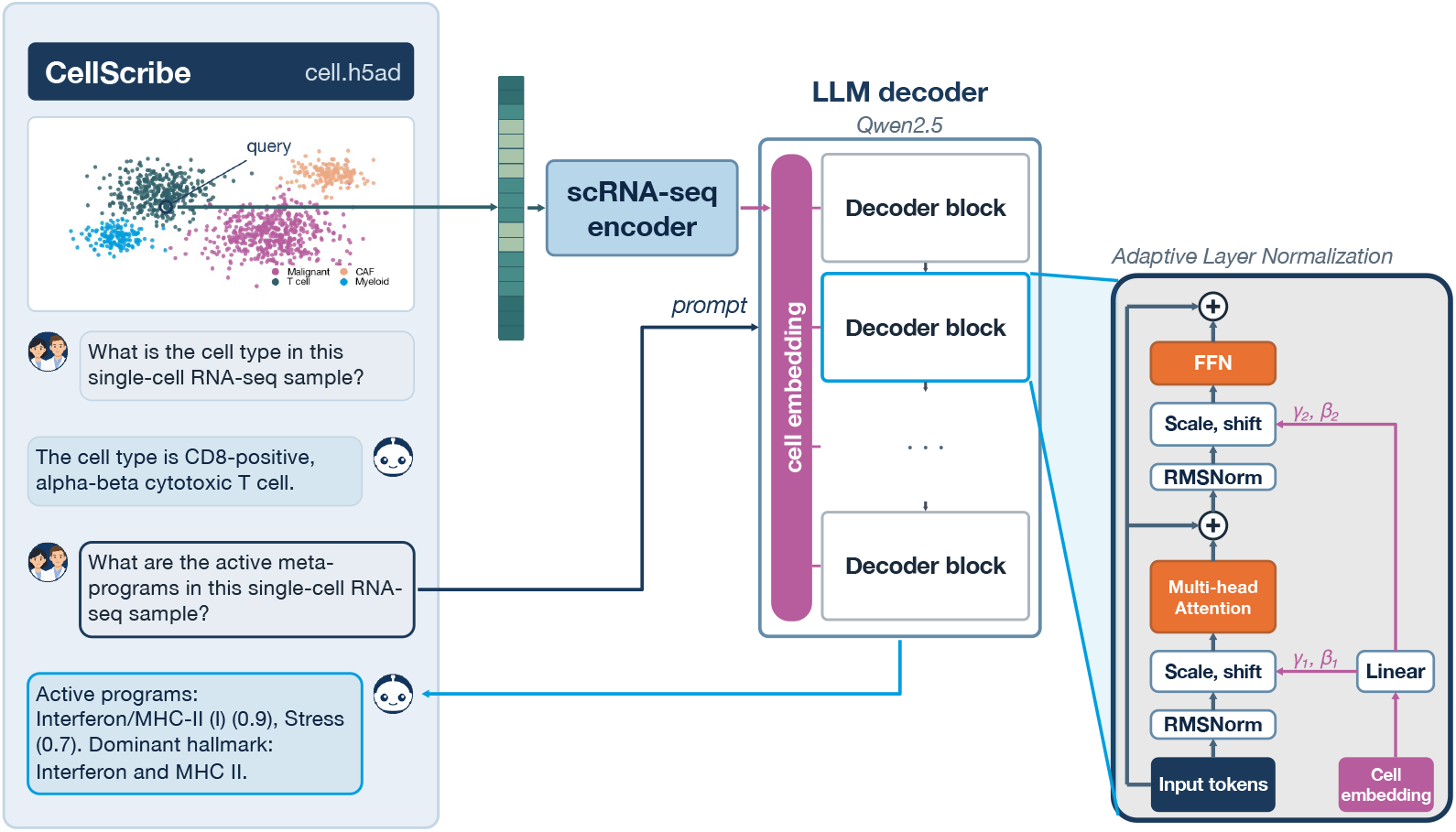
CellScribe architecture. A single-cell RNA-seq foundation model maps the expression vector to an embedding ***c***, while the task question is provided as tokens. Adaptive layer normalisation (AdaLN) conditions each decoder block on this embedding; the decoder generates either a cell type or a meta-program profile.

#### AdaLN-Zero conditioning

Rather than inserting the cell embedding as a special additional token in the LLM vocabulary, we use it to modulate the residual stream throughout the decoder. Our design is inspired by the AdaLN mechanism used in Diffusion Transformers [Peebles and Xie, 2023]. In block *l* and stream *s* ∈{attn, mlp}, a zero-initialised linear projection maps the 896-dimensional conditioning embedding directly to two vectors of the same dimension: the scale ***γ*** and shift ***β***. Formally, 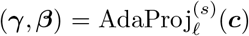. These vectors modulate the layer-normalised input [Ba et al., 2016]:

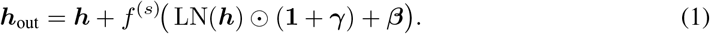

Zero-initialising the output projection gives ***γ***=***β***=**0** initially, so Eq. (1) recovers the pretrained sub-layer. During training, the single-cell encoder and LLM backbone remain frozen. Training updates only the projection that maps the cell representation into the LLM hidden space and the linear layers that predict the adaptive scale and shift. Algorithm 1 in Appendix B specifies the modulation as pseudocode, and the same appendix gives the optimisation settings.

### 3.2 Tasks, data and evaluation

#### Tasks

Cell typing and state prediction share the decoder and AdaLN pathway, but use different prompts and answer templates. Typing returns `“The cell type is {type}.”`; state uses a compact multi-program response: `“Active programs: Cell Cycle - G2/M (1.9), EMT-I (1.1), Stress (0.7). Dominant hallmark: Cell cycle.”` We emit at most *K*=5 canonical program names above activity 0.5, with one-decimal NMF magnitudes and a dominant hallmark (Appendix C).

#### Training corpora

CELLxGENE [CZI Cell Science Program et al., 2025] contributes 62.6M cells and more than 700 cell ontology types for cell typing. The 3CA corpus [Gavish et al., 2023] contains 4.07M cells from 99 cancer studies and provides activity scores for 41 meta-programs. We train state prediction on every 3CA cell with available scores, including cells from the tumour microenvironment rather than malignant cells alone. For typing, we refine the original “Malignant” annotation by tissue of origin, producing labels such as `malignant cell (Breast)` across 15 tissues. In studies shared by 3CA and CELLxGENE (500k cells), we match cell barcodes and transfer the corresponding fine-grained cell ontology labels to non-malignant cells, while preserving the tissue-refined labels of malignant cells. We remove the overlapping CELLxGENE datasets from training and use study-holdout splits throughout (Appendix A).

#### Evaluation

Our primary typing benchmark contains 4.44M cells from held-out CELLxGENE studies and 309 cell types; 50 additional cell types reserved for the unseen-type experiment (§6) are excluded. A prediction counts as correct only when it names the reference type. We report macro-*F*_1_, which weighs every type equally, and weighted-*F*_1_, which reflects type frequency. Hierarchical *F*_1_ extends the same comparison to the cell ontology: for a tolerance of *e* edges, we augment the predicted and reference types with their ancestors up to *e* levels and compute the *F*_1_ overlap of these sets. *hF*_1_@0 coincides with macro-*F*_1_; *hF*_1_@1 and *hF*_1_@2 give partial credit to nearby lineages. State metrics mirror the two-level organisation of the cell state. Dominant hallmark accuracy evaluates the coarse state; at the meta-program level, top-1 accuracy identifies the strongest program and set-*F*_1_ measures overlap among active programs. NDCG@5 [Järvelin and Kekäläinen, 2002] evaluates whether highly active programs are correctly retrieved and ranked, whereas mean absolute error (MAE) measures score error among programs present in both the prediction and reference. Appendices C and D provide formal definitions and baseline protocols.

## 4 Learning to Identify Cell Types and Quantify State

CellScribe is trained in three stages. The first, supervised fine-tuning, establishes that one language interface can perform both tasks; the two that follow exploit capabilities that fixed prediction heads do not naturally provide: post-training alignment of generations with biological rewards (§5) and in-context learning from labelled demonstrations to predict cell types absent from supervised training (§6). The benchmarks below report the complete three-stage model against the baselines, together with the score of supervised fine-tuning on its own; Appendices E and F report every stage separately.

The first stage trains the model jointly on cell typing and state prediction for 20M sampled training cells. For each cell, the model receives the corresponding task prompt and is trained by teacher forcing to generate the reference answer. Specifically, we minimise the autoregressive negative log-likelihood over answer tokens,

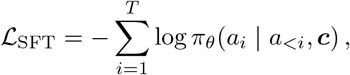

where *a*_*<i*_ comprises the task question and preceding reference answer tokens. A balanced sampler interleaves examples from the two tasks, so the same conditioning pathway learns both outputs without task-specific prediction heads. The objective updates only the cell projection and AdaLN scale-and-shift layers described in §3.

### Cell typing at scale

Figure 2a compares cell typing on the 309-type, 4.44M-cell study-holdout split. At this label-space scale, language and retrieval baselines perform poorly: Cell2Sentence, CellWhisperer and InstructCell reach macro-*F*_1_ scores of 0.083, 0.053 and 0.009, respectively. By contrast, a supervised probe of frozen scFoundation embeddings provides a strong discriminative reference, with macro-*F*_1_ 0.194 and weighted-*F*_1_ 0.366. Supervised fine-tuning of CellScribe alone respectively reaches 0.195 and 0.497, already matching the probe on the tail-sensitive metric while improving weighted-*F*_1_ by 13 points. The alignment and in-context stages add 2.8 macro-*F*_1_ points on top, so CellScribe reaches 0.223 and 0.484, ahead of the probe on both. The gain is concentrated in the tail: alignment redistributes recall from frequent types towards rare ones, trading weighted-*F*_1_ (0.497 to 0.479) for macro-*F*_1_, and the in-context stage recovers the weighted-*F*_1_ (0.484) while adding a further 1.8 macro-*F*_1_ points.

**Figure 2:**
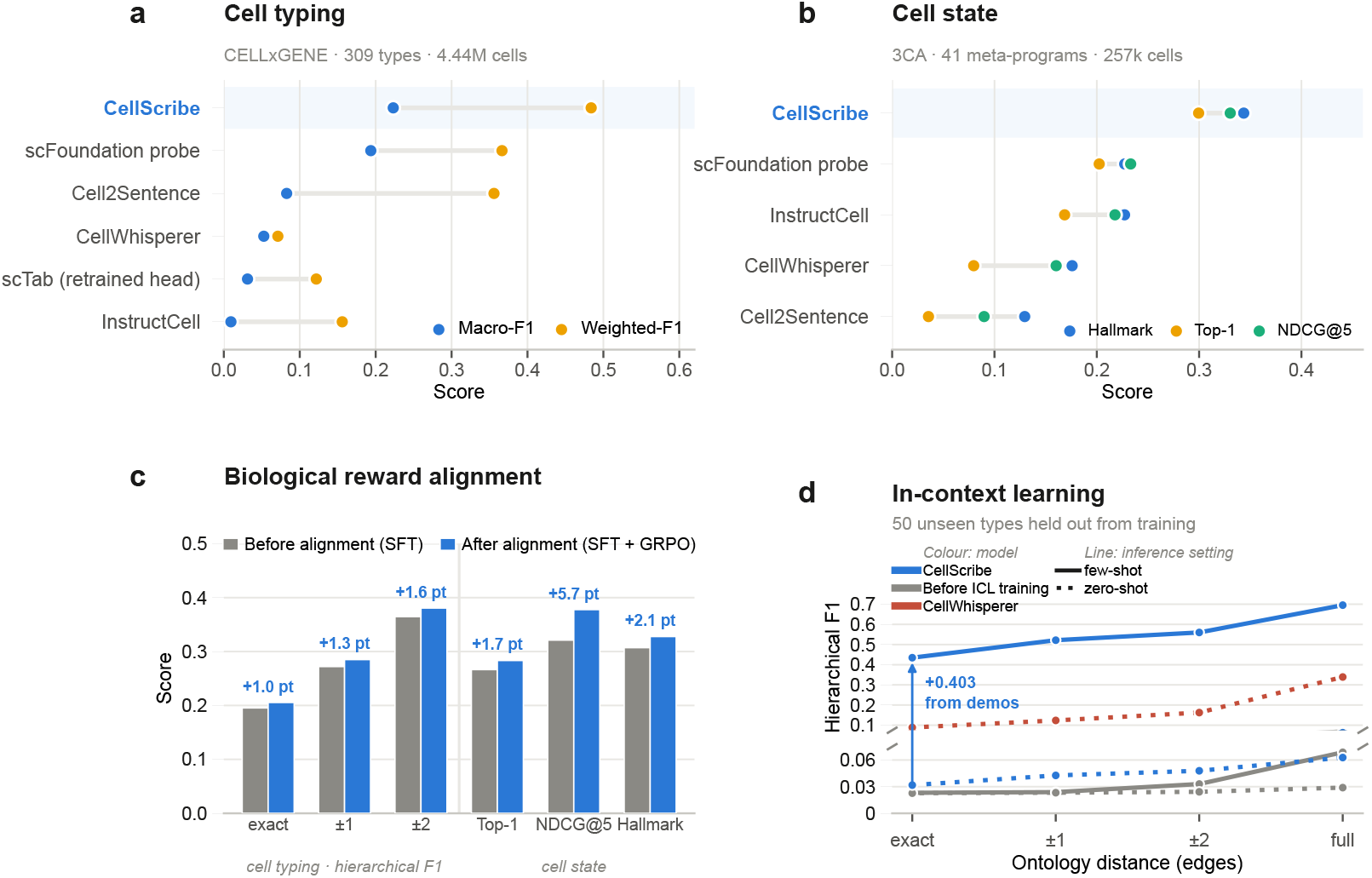
CellScribe evaluation. **(a)** CELLxGENE typing across 309 types and 4.44M cells, measured by macro- and weighted-*F*_1_. **(b)** 3CA state prediction across 41 meta-programs and 257k cells. **(c)** Post-training alignment with biologically grounded rewards improves hierarchical typing and state prediction. **(d)** Prediction of 50 cell types excluded from supervised training through span-AdaLN in-context learning with labelled cells. CellScribe always denotes the full pipeline, reported in **(a)** and **(b)**; **(c)** and **(d)** resolve it into stages, the supervised base before and after alignment in **(c)** and CellScribe against itself without the in-context post-training in **(d)**. Together the panels show that CellScribe combines large-vocabulary typing with state prediction and supports both biologically grounded alignment and demonstration-conditioned adaptation.

### Cell state quantification

State prediction is structurally more demanding than assigning one lineage: a cell can express several overlapping programs, whose membership, ranking and activity must be recovered jointly. On malignant cells from held-out 3CA studies (Figure 2b; Appendix F), the full three-stage CellScribe pipeline outperforms all baselines across these complementary criteria. It reaches 0.344 hallmark accuracy, 0.298 program-set *F*_1_ and 0.330 NDCG@5, compared with the probing baseline values of 0.227, 0.205 and 0.233, respectively; supervised fine-tuning alone already reaches 0.307, 0.270 and 0.321 (Appendix F). Thus, conditioning the LLM on a single-cell embedding supports a structured profile of co-active transcriptional programs rather than only a lineage label, and the model transfers this profile to unseen studies without cohort-specific NMF refitting.

### Focus on Tabula Sapiens

Tabula Sapiens [The Tabula Sapiens Consortium, 2022] is a widely used multi-organ reference atlas and one of the studies held out in our CELLxGENE test set. We use its 1.14M cells for a focused comparison with CellWhisperer (Figure 3). Across the full atlas and label space, CellScribe reaches 0.441 weighted-*F*_1_, versus 0.072 for CellWhisperer. We then examine the 16 well-studied types used in the CellWhisperer benchmark, including immune distinctions such as classical versus non-classical monocytes, macrophages versus monocytes, and CD4 versus CD8 T cells. On these 457k cells, CellScribe reaches 0.713 weighted-*F*_1_ over the full label space, compared with 0.473 for CellWhisperer. Restricting both models to the sixteen candidate types puts them in the same decision space, with CellScribe constraining its decoding to the sixteen candidate names and CellWhisperer retrieving against the same set (Appendix E.4), and CellScribe again leads, at 0.780 versus 0.683 weighted-*F*_1_. Table 9 in Appendix E.4 reports the complete metrics across all three conditions per model.

**Figure 3:**
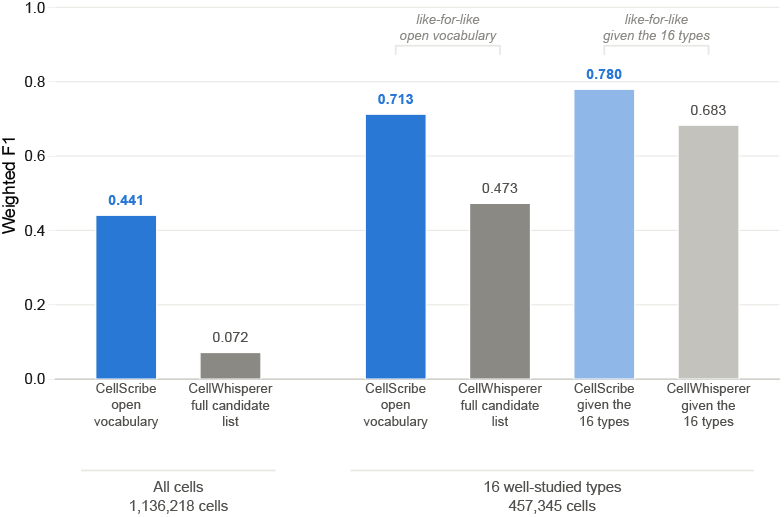
Tabula Sapiens typing (weighted-*F*_1_). The full-atlas analysis evaluates 1.14M cells over the complete label space; the focused analysis covers 457k cells from a subset of 16 common and immune-related cell types.

### Conditioning ablation

Figure 4 compares AdaLN with two alternative ways of conditioning the LLM on a cell embedding (Appendix G for exact figures). The `<scRNA-seq>` token approach projects the embedding into the LLM hidden space and inserts it as one prompt token. We test a single linear projection and a deeper multilayer projector sized to give the token model approximately the same number of trainable parameters as AdaLN. The Perceiver [Jaegle et al., 2021], in the resampler form used by Flamingo [Alayrac et al., 2022], uses learned latent queries to turn the cell embedding into 16 prompt tokens. AdaLN performs best on both tasks. Relative to AdaLN, the simple token projection, the parameter-matched token projection and the Perceiver reduce typing macro-*F*_1_ by 43%, 20% and 11%, respectively. The advantage is largest for cell state, where top-1 and hallmark accuracy fall by 50–80% under the alternative conditioning mechanisms. This ablation is run at the supervised fine-tuning stage.

**Figure 4:**
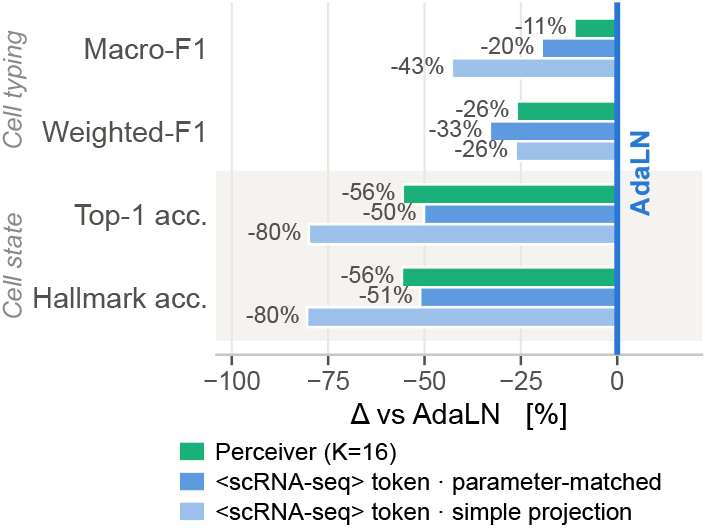
Conditioning ablations. Values are the results of alternatives to AdaLN on CELLxGENE typing and 3CA state prediction. Perceiver and token-projection alternatives reduce performance on both tasks.

## 5 Alignment with Biologically Grounded Rewards

### Motivation

Supervised likelihood teaches the model to reproduce reference answers, but treats each answer primarily as a token sequence rather than as a biological object. Cell identities instead form a graph in which some errors remain within the correct lineage, while cell state is a set of programs with graded activities and rankings. Because these outputs support biomedical interpretation, their error structure matters: confusing closely related immune subtypes differs from assigning the wrong lineage, while missing a dominant hypoxia or stress program can obscure tumour state. As CellScribe produces structured text, we can score each generated answer against these biological relationships and align the model after SFT using task-specific rewards.

#### Method

Starting from the SFT checkpoint, the selected alignment run processes 600k sampled training cells. Group Relative Policy Optimisation (GRPO) [Shao et al., 2024] samples *G*=12 completions per prompt. Standardised within-group advantages *Â* ^(*g*)^ = (*r*^(*g*)^ *−µ*)*/σ* drive a PPO-clipped update [Schulman et al., 2017], avoiding a value model. Typing reward is *R*_type_ = exp(*−d*_*G*_(*ŷ, y*)), where *d*_*G*_ is cell ontology shortest-path distance, plus an out-of-vocabulary penalty. State reward mixes program-set agreement, score error, hallmark agreement and parseability:

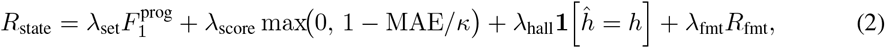

where MAE is computed over programs shared by predicted and reference sets, *h* is the dominant hallmark, and *R*_fmt_ scores whether the generation parses. Appendix H specifies both rewards in full, including the component weights.

#### Alignment results

Grounding generation in the ontology reward sharpens cell typing where it matters: residual mistakes move closer along the cell ontology. Hierarchical *F*_1_ rises by 1.0, 1.3 and 1.6 points at exact, *±*1 and *±*2 tolerance (Figure 2c), and the lift growing with tolerance shows that the remaining errors are pulled onto nearer ontology neighbours rather than merely converted into additional exact hits, exactly the hierarchical behaviour the reward is designed to induce. Cell state gains alongside typing, with top-1 program accuracy, hallmark accuracy and NDCG@5 up by 1.7, 2.1 and 5.7 points, the last indicating markedly better ranking of the active programs.

#### Reward granularity and skipped groups

The two tasks are scored at very different resolutions. Cell typing reward is a function of a single integer ontology distance and takes on the order of ten values, whereas cell state combines four partially independent terms over a scored set of 41 programs and ties only when all four agree at once. Because GRPO standardises rewards within a group of *G*=12 completions, a group whose rewards are all equal produces no gradient and is discarded, and the coarser typing reward loses far more groups: 26.6% against 5.0% for cell state in our GRPO model. These ties are exact rather than merely close, and in roughly 80% of them the twelve completions are identical and already correct, so no reward-side change recovers them. We therefore adopt DAPO-style dynamic sampling [Yu et al., 2025]: each step oversamples candidate cells threefold, screens each with two cheap generations, and spends the full rollout only on the most contested. In a single-factor comparison against the same run without this prescreening (Appendix H.2), this roughly halves the mean typing skip rate, 48.0% to 26.6%, at 1.5*×* the generation cost.

## 6 Towards In-Context Adaptation to Unseen Cell Types

### 6.1 Conditioning several cells in one prompt

#### Motivation

A closed-set classifier cannot predict a type absent from its output head. A generative model can instead compare a query with labelled cells in its prompt, but this requires extending AdaLN to condition one prompt on multiple cells while preserving which token span corresponds to each cell.

#### Span-conditioned AdaLN

Standard AdaLN associates one cell embedding with the entire token sequence, so it cannot distinguish several cells placed in the same prompt. We replace this global assignment with a position-dependent one (Figure 5). Each demonstration cell and the query cell are first encoded independently, giving ***c***_*i*_ = *E*(***x***^(*i*)^) for demonstration *i* and ***c***_*q*_ = *E*(***x***^(*q*)^) for the query. We then associate every token position with the cell it describes: tokens in demonstration *i* use ***c***_*i*_, whereas the system, query and answer tokens use ***c***_*q*_. Formally, for *M* demonstrations, *π* : {1, …, *T }→ {q*, 1, …, *M}* records this assignment. In block *l* and stream *s* ∈{attn, mlp}, token *p* is modulated by

**Figure 5:**
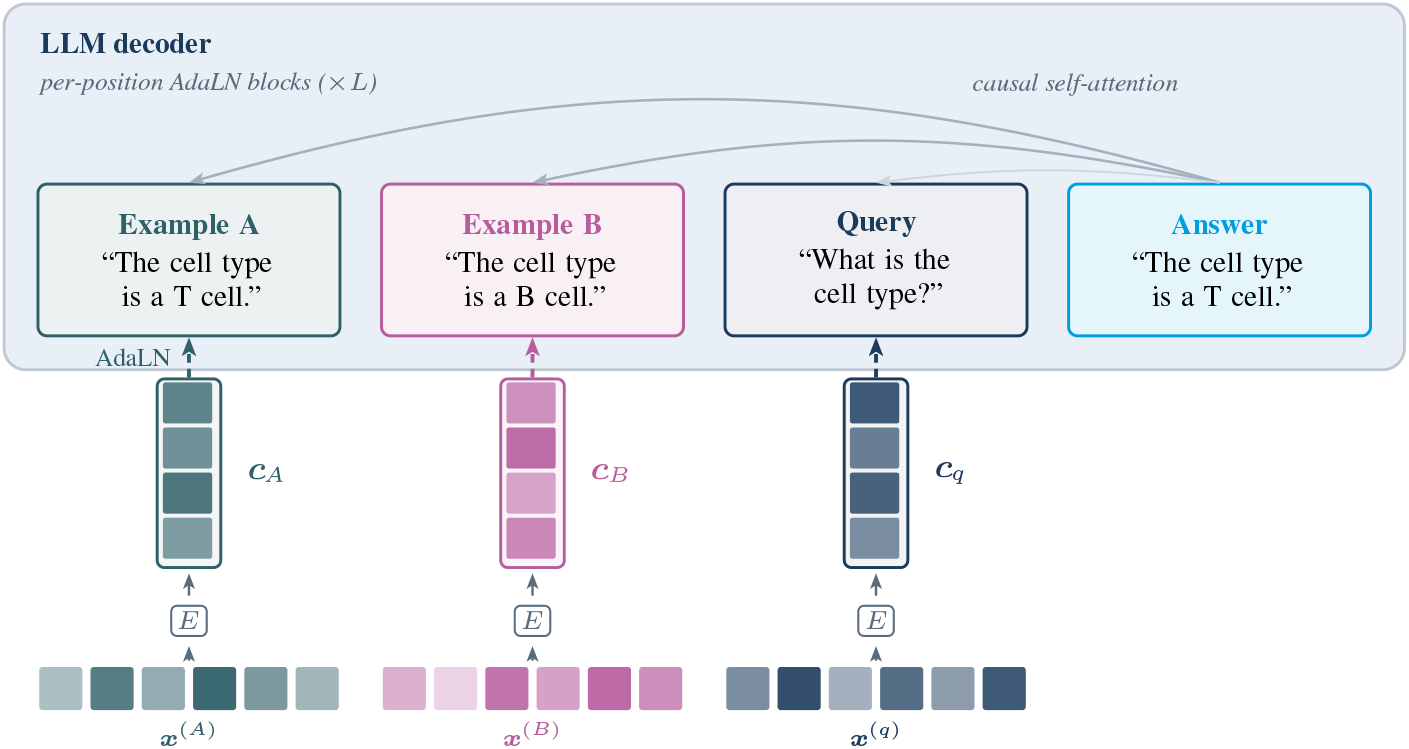
Span AdaLN (Eq. (3)). Each demonstration modulates its decoder-token span with its own cell embedding; the query span uses the query embedding. Causal self-attention conveys labelled supports to the answer.

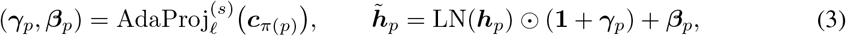

and, writing ***H*** = (***h***_1_, …, ***h***_*T*_ ) and 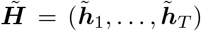 for the token vectors stacked over the sequence, the sub-layer acts as 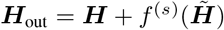. Causal self-attention then lets the answer use the labelled demonstrations. With no demonstrations, every position uses ***c***_*q*_, recovering standard AdaLN in Eq. (1).

We add a third training stage on top of the two above: starting from the aligned checkpoint of §5, that is the SFT model after GRPO, we continue training for 6M cells on multi-cell prompts interleaved with ordinary single-cell batches; more details on this training stage can be found in Appendix I.

### 6.2 Evaluating on unseen types

#### Protocol

We hold out 50 cell types from all three training stages: supervised fine-tuning, reward alignment and the multi-cell post-training above. We sample their cells from the test set. Many are fine-grained composite ontology terms that combine lineage, anatomy and subtype, such as “kidney loop of Henle thick ascending limb epithelial cell”. For each held-out type we split its test cells into a demonstration pool of at most 64 cells and a disjoint query pool of at most 100, so no cell is ever both, which yields 4,671 query cells in total. Each query defines a three-way, one-shot episode: the prompt contains one labelled support cell from the query’s type and one from each of two randomly sampled held-out distractor types. The correct type is therefore absent from supervised targets but demonstrated once in context. We use the same episodes across models and ablations.

#### Surrogate labels

Post-training episodes must use cell types seen during SFT so that the 50 evaluation types remain genuinely unseen. With their real names, however, the model can predict the familiar label directly from the query embedding and ignore the supports, a form of shortcut learning [Geirhos et al., 2020]. Following symbol tuning [Wei et al., 2023], we replace type names with a new set of ontology-shaped nonce phrases in every episode. Because the query embedding has no fixed association with these arbitrary labels, the model must match the query to a demonstration and copy the label attached to that cell. At evaluation, the supports again use real cell ontology terms, testing whether this learned binding mechanism transfers to unseen names.

#### In-context discovery of held-out cell types

Figure 2d first evaluates whether labelled cells enable the model to name unseen types without restricting its output vocabulary. The post-trained arm is CellScribe itself, and the base arm is the aligned checkpoint of §5, which is the same model with this post-training removed, so the comparison isolates the post-training. With the correct demonstrations, that post-training raises exact hierarchical *F*_1_ from 0.023 to 0.435; the gain persists at every ontology tolerance, reaching 0.695 over the full lineage against 0.069 for the base given the same demonstrations. Removing the demonstrations eliminates the gain (0.032 exact), confirming that the model relies on the in-context cell label bindings rather than a memorised type prior. The post-trained model also exceeds the CellWhisperer zero-shot reference across the complete hierarchical curve, demonstrating the benefit of labelled cells over zero-shot retrieval.

#### Disentangling generation reveals stronger unseen-type identification

Free generation asks two things at once: identify the matching demonstration, then reproduce a term the model was never trained to emit. Retrieval baselines such as CellWhisperer are asked only the first, as they score a supplied vocabulary. We therefore add two candidate-aware readouts (Figure 6, with exact numbers in Table 14 of Appendix I). The first restricts decoding to a candidate set, either the full vocabulary, the 50 held-out types or the three demonstrated types. The second scores every one of the 50 held-out names as a continuation of the prompt and predicts the one with the highest length-normalised log-likelihood (Eq. (4)). Exact match rises from 0.364 unrestricted to 0.525, 0.819 and 0.882 as the candidate set narrows; ranking those same 50 names by likelihood instead of decoding them reaches 0.433. This ability comes from the demonstrations rather than the candidate set: permuting the support labels lowers the three constrained scores to 0.091, 0.245 and 0.302 and likelihood ranking to 0.163, and removing the supports lowers likelihood ranking further, to 0.091. The model without this post-training, in contrast, scores the same whether the support labels are correct or permuted, at every candidate set. The ICL post-training is therefore what makes the model read the support cells and answer with the label bound to the one that matches the query. Surrogate labels create this sensitivity: training the same recipe from the same checkpoint on real names drops free generation to 0.040, full-vocabulary accuracy to 0.090 and likelihood rank 1 to 0.135.

**Figure 6:**
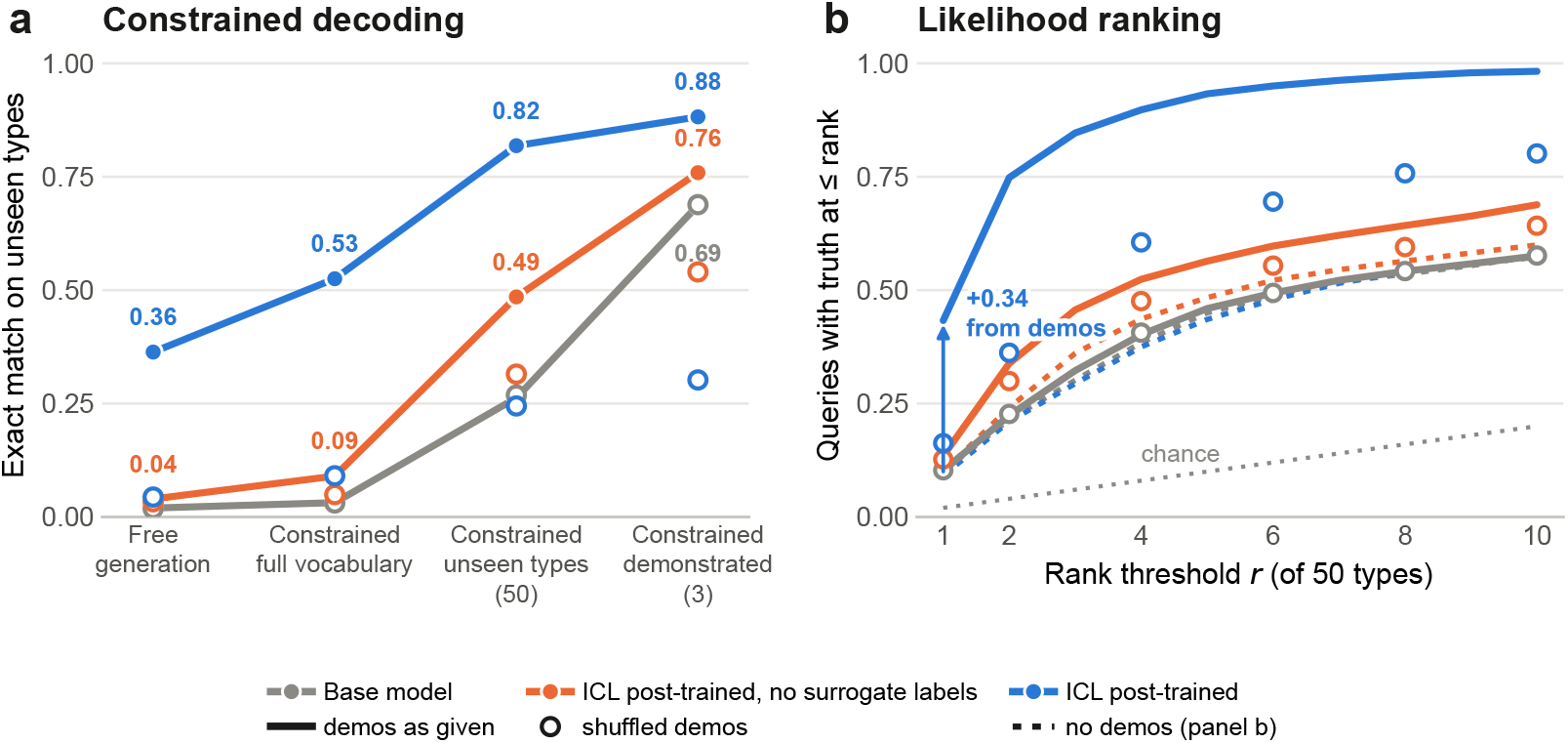
In-context classification of unseen types. **(a)** Greedy decoding when unrestricted, or constrained to the full vocabulary, the 50 unseen types or the 3 demonstrated types. **(b)** Likelihood ranking over the 50 held-out types. Filled marks use correct supports, hollow marks shuffle their labels, and the dotted curves in **(b)** omit the supports entirely. Results cover 4,671 queries.

## 7 Conclusion

We presented CellScribe, an AdaLN-conditioned LLM that casts cell typing and cell state prediction as a single generative task, mapping one expression profile to a fine-grained ontology type or to a ranked set of scored meta-programs through the same language interface. It achieves the strongest overall cell typing results in our benchmark and is especially strong on cell state. This second task is more demanding than assigning one label: the model must recover several co-active programs, rank them and estimate their activity. The language interface directly enables two additional capacities: aligning generated answers with biological knowledge and learning from labelled cells in context. GRPO optimises ontology distance and meta-program agreement directly, improving hierarchical typing and state prediction over SFT. Span-conditioned post-training teaches the model to read several labelled cells in one prompt and identify types entirely excluded from supervised training, extending annotation from a fixed vocabulary to demonstration-based adaptation. A dataset containing types that no training vocabulary covers can then be annotated from a handful of labelled cells, without retraining the model or refitting a classifier head, which is precisely where closed-set annotators fail. Together, these results show that one conditioned LLM can combine fine-grained cell typing, quantitative state profiling, biological alignment, and in-context discovery.

### Cross-study cancer profiling

One health application motivates these capabilities: research-grade profiling of tumour ecosystems in new cohorts. CellScribe scores program activity per cell without refitting NMF on each cohort, and our study-holdout evaluation measures exactly that transfer. Because it retains co-active proliferation, hypoxia, stress, and epithelial-to-mesenchymal transition, instead of imposing one state label, its cell-resolved output can be tested directly against clinical biomarkers, treatment response, and patient outcomes in prospective cohorts.

### Limitations and open questions

The strong performance of probes built on pretrained scFoundation embeddings shows that these encoders already contain highly discriminative structure for cell typing. Cell state is a more demanding structured prediction problem, and the particularly strong results suggest that LLM-based models are well-suited to represent overlapping, quantitative biological processes. Perturbation prediction is a natural next step: conditioning jointly on a cell and a drug or genetic intervention could predict which programs change and describe the resulting state transition, and would test whether this capacity extends beyond the current 41 NMF-derived meta-programs. A further interpretability question is how AdaLN uses the cell embedding throughout generation: how its scale and shift modulations act across decoder layers and at each decoding step to produce type and state tokens. In parallel, ongoing work couples bulk RNA-seq deconvolution to CellScribe, so that the cell-type profiles recovered from a bulk sample can be typed and scored by the model, linking its outputs to clinical cohorts and endpoints. That integration falls beyond the present paper, whose goal is to introduce and validate the method on single-cell data, and will require prospective patient-level evaluation.

## A Data, harmonisation and evaluation splits

### Source datasets

CELLxGENE Census v2 [CZI Cell Science Program et al., 2025] (2025-01-30 snapshot) contains 62.6M cells from 479 datasets, 144 tissues and 794 cell ontology types. The Curated Cancer Cell Atlas (3CA) [Gavish et al., 2023] contains 4.07M cells from 99 cancer studies and scores for 41 meta-programs, grouped into the 11 hallmarks in Table 1.

**Table 1:**
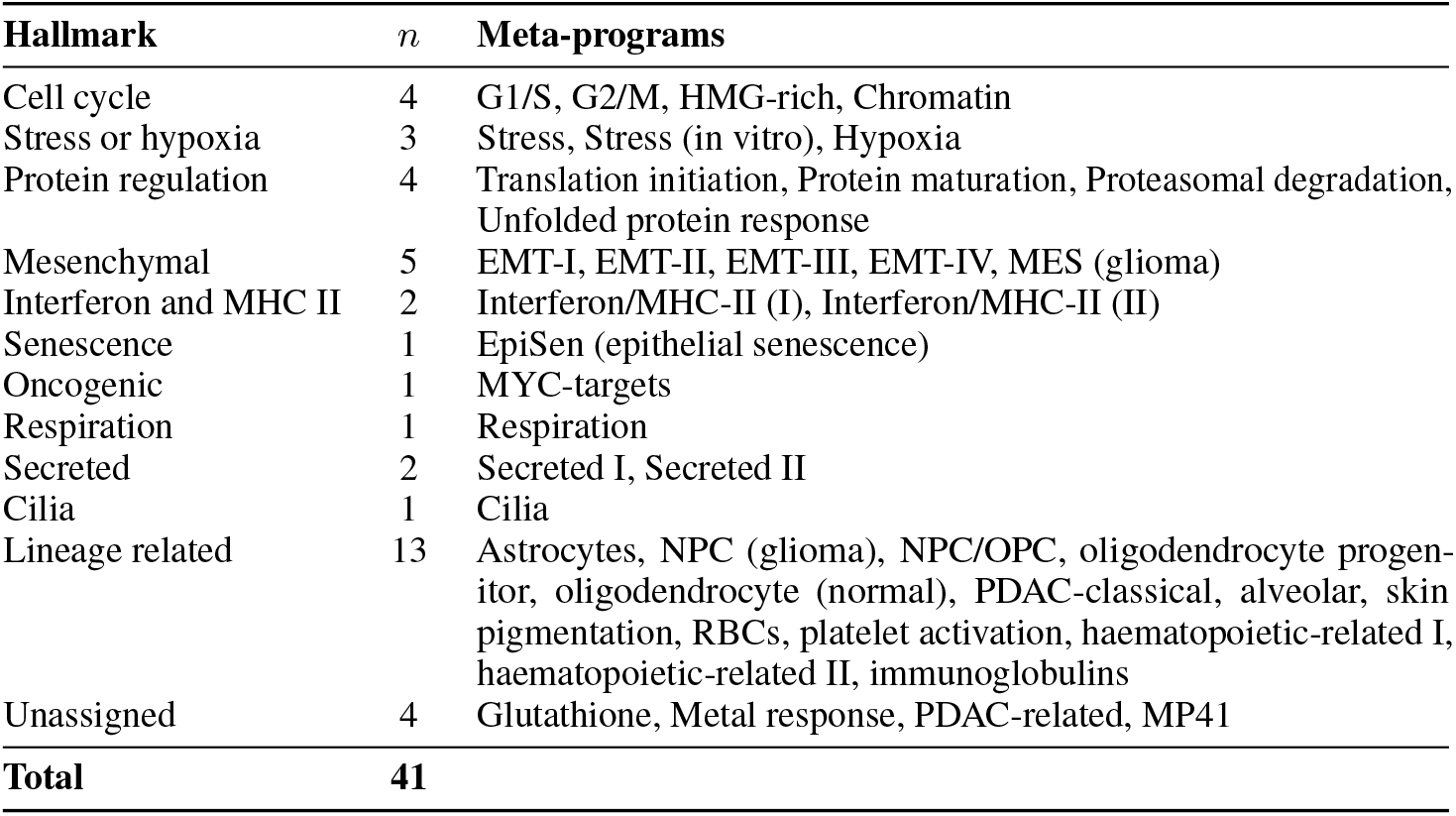
The 41 consensus malignant meta-programs organised into 11 hallmarks plus an Unassigned category, following Gavish et al. [2023].

### Merging the two label sets

We map 66 of the 79 abbreviated 3CA labels to the cell ontology, covering 98.5% of typed 3CA cells. Six 3CA studies (500k cells) overlap with CELLxGENE, as verified by metadata and cell barcodes. We exclude the corresponding 10 CELLxGENE datasets from training and transfer their fine-grained labels only to non-malignant 3CA cells, as 3CA annotations tend to be coarser than those available in CELLxGENE. The 3CA “Malignant” label is retained because CELLxGENE does not encode malignancy consistently. We add 15 tissue-specific malignant labels, such as `malignant cell (Breast)`, to the evaluation ontology.

### Study-holdout training and test splits

All partitions are study-holdout: every study belongs to a single partition, so no test cell shares a batch, protocol or donor with a training cell. The primary typing split excludes the 50 types reserved for unseen-type evaluation (§6); Table 2 gives the resulting cohorts.

**Table 2:** Evaluation cohorts, grouped by task. All are drawn from the held-out studies and are selected by different criteria, so they are not disjoint: the two Tabula Sapiens rows are a single CELLxGENE test study whose cells are also counted in the primary typing benchmark, and the held-out-type cohort is selected by label rather than by study. “Labels” counts cell types, except for cell state where it counts meta-programs; “Studies” counts the distinct source studies contributing cells.

| Evaluation cohort | Corpus | Labels | Studies | Cells |
| --- | --- | --- | --- | --- |
| *Cell typing* |  |  |  |  |
| Primary benchmark (reserved types excluded) | CELLxGENE | 309 | 48 | 4,444,795 |
| Held-out types (50 reserved types) | CELLxGENE | 50 | 36 | 1,904,139 |
| Tabula Sapiens, full atlas | CELLxGENE | 180 | 1 | 1,136,218 |
| Tabula Sapiens, focused subset | CELLxGENE | 16 | 1 | 457,345 |
| Cancer benchmark | 3CA | 40 | 16 | 661,147 |
| *Cell state* |  |  |  |  |
| Meta-program benchmark | 3CA | 41 | 16 | 256,978 |

## B Architecture and optimisation

This appendix gives the conditioning mechanism as pseudocode and the optimisation settings of the supervised and alignment stages.

### B.1 AdaLN-Zero conditioning

Each of the 24 decoder blocks has two AdaLN projectors, one per layer normalisation (Algorithm 1). A projector maps the 896-dimensional cell embedding to scale and shift vectors. Zero initialisation preserves the pretrained decoder at the start of training.

#### Algorithm 1

AdaLN-Zero conditioning for decoder block *l*

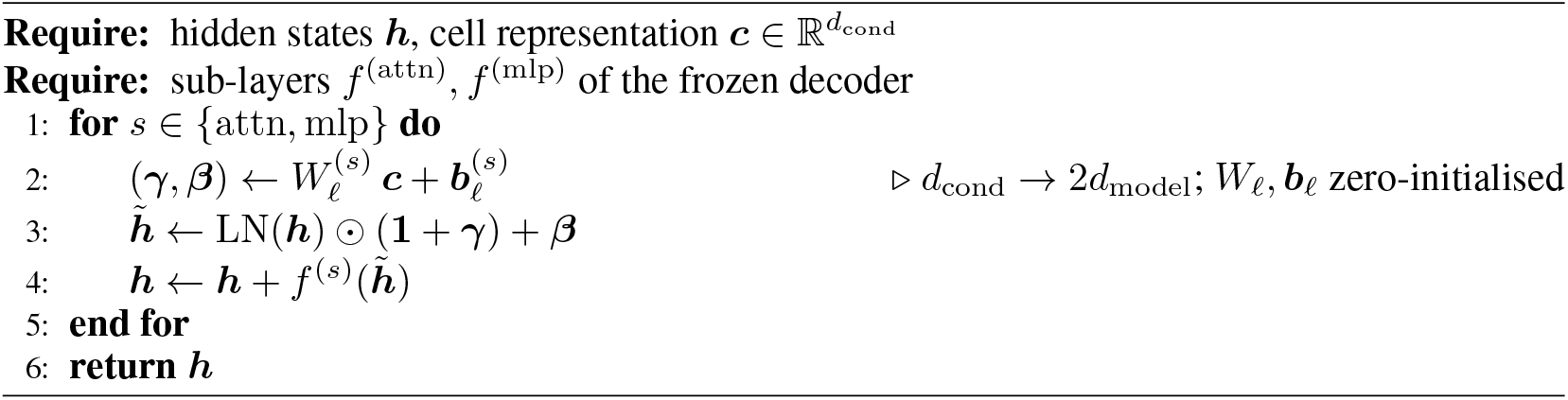

For span conditioning (§6) the only change is that ***c*** becomes position-dependent: line 2 is evaluated per token position *p* with ***c***_*π*(*p*)_, and lines 4–5 run over the whole sequence, as in Eq. (3).

### B.2 Supervised fine-tuning and alignment hyperparameters

#### Supervised fine-tuning

scFoundation and Qwen remain frozen, so the trainable parameters are the linear projection of the cell representation into the decoder hidden space and the 48 AdaLN projectors, 79.9M parameters in total. We optimise with AdamW [Loshchilov and Hutter, 2019] at a peak learning rate of 2*×*10^*−*4^ and weight decay 0.01, after 1000 warm-up steps, followed by a cosine decay towards 2*×*10^*−*5^. The effective batch is 64 cells, and training runs in bfloat16 with gradient-norm clipping.

#### Reward alignment

GRPO starts from the supervised checkpoint and processes 600k cells. Each prompt is answered by *G*=12 completions sampled at temperature 1.3; rewards are standardised within the group, the resulting advantages are clipped to *±*3, and groups whose reward standard deviation falls below 0.01 are skipped. The effective batch size is 8, and the optimiser is again AdamW, at 5*×*10^*−*6^ with weight decay 0.01, 100 warm-up steps and a cosine decay to 5*×*10^*−*7^. Alignment updates the same parameters as supervised fine-tuning, the cell projection and the AdaLN projectors.

## C Prompts, targets and metrics

### C.1 Prompts and answer templates

Table 3 gives the shared system prompt and task-specific templates.

**Table 3:** Prompts and answer templates. The system prompt is shared by both tasks; the user prompt and the answer template are task-specific.

|  |  |  |
| --- | --- | --- |
| **System prompt** *shared by both tasks* |  |  |
| You are a helpful assistant who answers questions about cell types and active transcriptional meta-programs from single-cell RNA-seq data. |  |  |
|  | **Cell typing** | **Cell state** |
| User prompt | What is the cell type in this single-cell RNA-seq sample? | What are the active meta-programs in this single-cell RNA-seq sample? |
| Answer template | The cell type is {type}. | Active programs: {program} ({score}), {program} ({score}). Dominant hallmark: {hallmark}. |

State targets retain the top *K*=5 programs whose NMF activity exceeds a threshold *τ* =0.5, ordered by activity and rounded to one decimal. The dominant hallmark follows the highest-scoring program in the 41-program registry of Gavish et al. [2023], grouped into the 11 hallmarks in Table 1. If none crosses the threshold, the response reports only the highest-scoring program and its hallmark.

### C.2 Answer parsing and likelihood scoring

For unconstrained typing, we extract the predicted type with a simple string-based parser; unparsed responses are recorded as `Unknown`. Trie-constrained decoding, used for the constrained readouts of §6, instead restricts generation at every step to the tokens that extend a prefix of some vocabulary term, so its outputs are valid terms by construction and need no parsing. For cell state, a parser extracts the reported programs, scores and hallmark.

Alongside generation, free or constrained, we also use a likelihood-based readout, which scores a fixed set of candidate strings instead of generating an answer. Given a prompt *P*, a conditioning vector ***c*** and a candidate *y* tokenised as (*y*_1_, …, *y* |_*y*_|), we take the length-normalised mean token log-likelihood

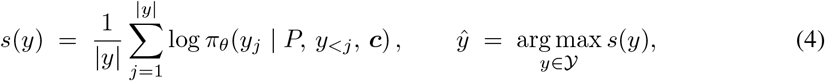

over the candidate set *Y*. Length normalisation limits the preference for short names. This requires |*Y* | forward passes per cell and yields a full candidate ranking. We use Eq. (4) for the CellScribe likelihood ranking of §6 and for the generative baselines of Appendix D, with *Y* the 50 reserved type names and the 41 meta-program names respectively.

### C.3 Evaluation metrics

#### Metrics for cell typing

We use two families of metric.

- **Exact match**. Weighted-*F*_1_, macro-*F*_1_, weighted precision and weighted recall over the predicted and reference type labels, all requiring the exact term.
- **Hierarchical** *F*_1_. A prediction is credited by how much of the cell ontology lineage it shares with the target. Let An_*e*_(*v*) denote *v* together with the ancestors lying within *e* edges of it in the cell ontology; the “full lineage” setting drops the truncation and keeps every ancestor up to the root. A cell with target *y* and prediction *ŷ* contributes the overlap |An_*e*_(*ŷ*)∩An_*e*_(*y*) |, normalised by |An_*e*_(*ŷ*) |for hierarchical precision and by |An_*e*_(*y*)| for hierarchical recall. For each type, precision is accumulated over the cells predicted as that type and recall over the cells whose reference label is that type; the two are combined into *hF*_1_@*e* per type and averaged over types. At *e*=0 each ancestor set is the type itself and *hF*_1_@0 coincides with macro-*F*_1_, so increasing *e* relaxes the same metric rather than replacing it. `Unknown` predictions are scored at the graph root.

#### Metrics for cell state

Let *Ŝ* and *S* be the sets of programs named in the prediction and in the reference, each ordered by its reported activity. All five metrics are computed per cell and averaged over cells.

- **Top-1 program accuracy**. The fraction of cells whose highest-scoring predicted program is the reference’s most active one, so it scores the leading program only.
- **Dominant-hallmark accuracy**. The fraction of cells whose predicted dominant hallmark matches the reference one, a coarser readout over the 11 groups of Table 1.
- **Set precision, recall and** *F*_1_. Order and scores are ignored and the named programs alone are compared, as *P* = |*Ŝ ∩ S*|*/*|*Ŝ*|, *R* = |*Ŝ ∩ S*|*/*|*S*| and *F*_1_ = 2*PR/*(*P* + *R*).
- **NDCG@**5. The normalised discounted cumulative gain scores the order of the predicted programs, grading each position by the reference activity of the program placed there, so it rewards ranking the truly active programs first. Write *ŷ*_*i*_ for the program the prediction ranks at position *i*, and let the graded relevance rel(*p*) of a program *p* be its reference NMF activity in this cell, that is the value the reference answer would report for it, and 0 when the reference does not list it. A predicted program therefore contributes in proportion to how active it truly is, and an inactive one contributes nothing. With this relevance,

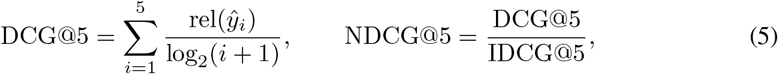

where IDCG@5 is the largest value the sum can take, reached by placing the five most active reference programs in decreasing order of activity. NDCG@5 is therefore 1 exactly when the prediction returns those five programs in that order.
- **Score mean absolute error**. The mean absolute error (MAE) between the predicted and reference activities, taken over *Ŝ* ∩ *S*, so the numeric scores are measured only for correctly named programs. It is undefined for the ranking-only baseline protocols of Appendix D, which return names without scores.

## D Baseline models and inference protocols

We evaluate scTab, CellWhisperer, Cell2Sentence and InstructCell on the same study-holdout splits, labels and metrics as CellScribe. Cell typing is native to all four: each was built and released to annotate cells, so we query each one the way its authors do and report that. We also describe, in §D.5, the probe on frozen scFoundation embeddings that serves as a discriminative reference on both tasks.

scTab is a closed-set classifier with no text interface: it selects an index in a fixed label head and cannot name anything outside it. We therefore reserve it for cell typing. The other three are text-based, and although none has been trained on the 41 meta-programs of Gavish et al. [2023], each has accumulated knowledge that can plausibly transfer to them, which is why we still consider them as cell-state baselines. CellWhisperer aligns expression with natural-language summaries that a language model writes from study and sample-level metadata, whose fields cover disease, treatment and experimental condition as well as tissue and cell type; its authors report that the resulting space captures cell types, tissues, diseases and developmental stages, and validate it by retrieving gene-set labels such as “response to type I interferon”. Cell2Sentence and InstructCell are generative models built on general-domain language-model backbones, Gemma 2 and T5 respectively, then tuned across single-cell tasks, cell-type and tissue annotation among them.

None of these three models exposes an interface for cell state, so every cell-state inference protocol below is one we constructed on their behalf. Rather than let a single arbitrary choice of ours stand for a model, we build several per baseline and report each at its best: four query phrasings for CellWhisperer’s retrieval, and both greedy generation and likelihood ranking (Eq. (4)) for the two generative models.

The protocols do not all support the same metrics. Greedy generation returns a single program name, so NDCG@5 has one ranked item to work with instead of five and makes considerably less sense on those rows than for the protocols that rank the whole program set; likelihood ranking, by contrast, orders all 41 names and is the protocol we report for the two generative models in the main text. Conversely, CellScribe is the only model here that reports an activity per program, so it is the only one with a defined score MAE: likelihood ranking orders the program names without scoring them, greedy generation returns a bare name, cosine similarities lie on an arbitrary scale, and the probe emits probabilities of activity rather than activities.

### D.1 scTab

scTab [Fischer et al., 2024] is a tabular cell-type classifier. Its released head covers 164 types and overlaps only partly with our test vocabulary. We therefore freeze its pretrained feature extractor and fit a new classification head on our training split over the full benchmark label space, and predictions are the arg-max of this head.

### D.2 CellWhisperer

CellWhisperer [Schaefer et al., 2025] is a CLIP-style dual encoder [Radford et al., 2021] (Geneformer transcriptome encoder, BioBERT text encoder [Lee et al., 2020]) that scores a cell against a text query by cosine similarity. A prediction is whichever candidate text in a supplied list scores highest, so that list defines its decision space. For cell typing we supply as candidates the cell types of our vocabulary, except in the Tabula Sapiens experiment of Table 9, where the candidates are restricted to the sixteen types of the focused subset.

For cell state, we rank the 41 program queries by cosine similarity; the first result determines top-1 and hallmark accuracy, and the first five determine set-*F*_1_ and NDCG@5. We report no score MAE for this model: a cosine similarity lives on an arbitrary scale and is not an activity value, so comparing it with a reference activity would not be meaningful. CellWhisperer defines no meta-program query, so we adapt the phrasing the authors use for their own zero-shot label retrieval, “Sample of a {label}”, to program names, and test three richer formulations against it (Table 4). Under the four formulations (Table 5), program names give the best hallmark accuracy, while the full descriptions of the meta-programs given by Gavish et al. [2023] give the best set-*F*_1_ and NDCG@5; naming the hallmark or appending marker genes helps neither. Program names are the phrasing we report for CellWhisperer both in the main text and in Table 10, as it stays closest to the authors’ own protocol.

**Table 4:**
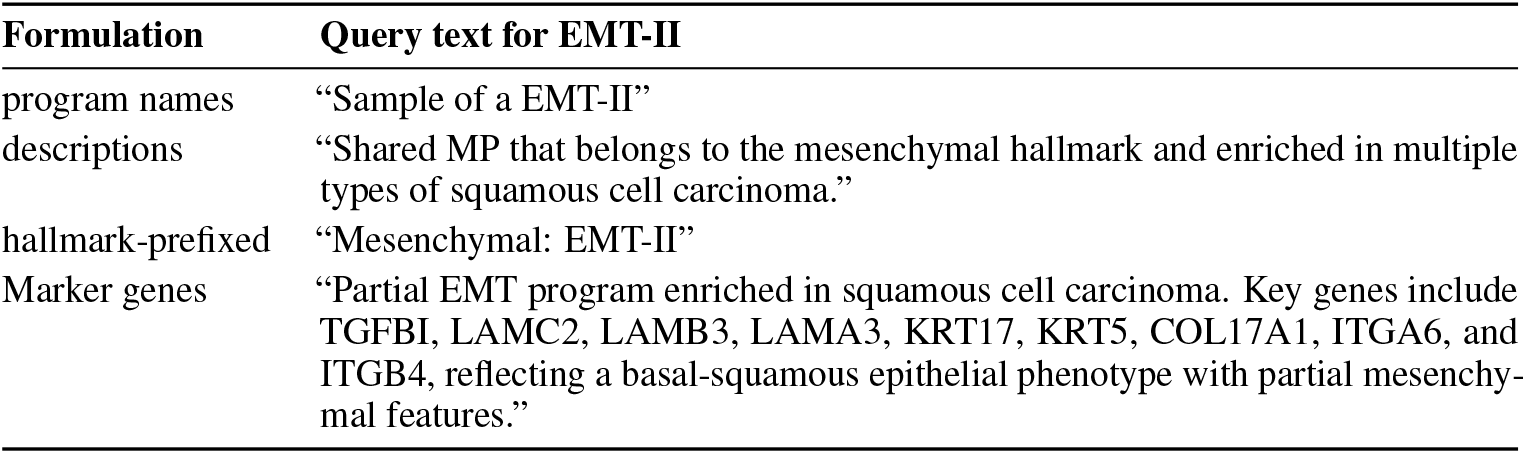
The four text queries, shown in full for the *EMT-II* program. Every formulation is applied to all 41 programs the same way. Program names and hallmark-prefixed names are built from the program’s label; the other two are the program’s description in Gavish et al. [2023], with marker genes appended in the last.

| Formulation | Query text for EMT-II |
| --- | --- |
| program names | “Sample of a EMT-II” |
| descriptions | “Shared MP that belongs to the mesenchymal hallmark and enriched in multiple types of squamous cell carcinoma.” |
| hallmark-prefixed | “Mesenchymal: EMT-II” |
| Marker genes | “Partial EMT program enriched in squamous cell carcinoma. Key genes include TGFBI, LAMC2, LAMB3, LAMA3, KRT17, KRT5, COL17A1, ITGA6, and ITGB4, reflecting a basal-squamous epithelial phenotype with partial mesenchymal features.” |

**Table 5:**
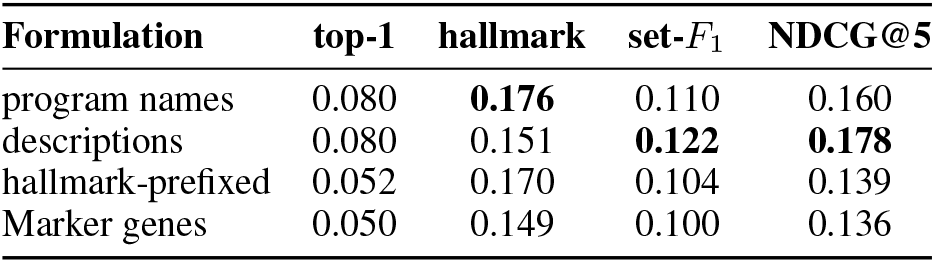
Cell-state performance of CellWhisperer under each of the four query formulations of Table 4.

### D.3 Cell2Sentence

Cell2Sentence (C2S-Scale-Gemma-2-2B [Levine et al., 2024, Gemma Team, 2024]) encodes a cell as a “cell sentence”, the names of its top 1000 genes ordered by decreasing expression, and generates text. Typing uses greedy generation followed by the same parser used for CellScribe, from the authors’ own prompt: *“The following is a list of* 1000 *gene names ordered by descending expression level in a Homo sapiens cell. Your task is to give the cell type which this cell belongs to based on its gene expression. Cell sentence: {sentence}. The cell type corresponding to these genes is:”*.

Cell state keeps the same construction with the question restated: *“The following is a list of* 1000 *gene names ordered by descending expression level in a malignant Homo sapiens cell. Your task is to identify which of the following malignant transcriptional meta-programs is active in this cell based on its gene expression. Possible meta-programs: {the* 41 *names}. Cell sentence: {sentence}. The meta-program corresponding to these genes is:”*. We compare two ways of turning this prompt into an answer: greedy generation and length-normalised likelihood ranking (Eq. (4)). Likelihood ranking is strongest, and is the C2S protocol we report in the main text and in Table 10.

### D.4 InstructCell

InstructCell [Fang et al., 2025] is a Q-Former [Li et al., 2023]+T5 [Raffel et al., 2020] model that consumes a gene-expression vector and a natural-language instruction. Typing uses greedy generation, from the instruction of the authors’ own demonstration: *“Can you help me annotate this single cell from a {species}? It was sequenced using {sequencing method} and is derived from {tissue}. The gene expression profile is {input}. Thanks!”*. Cell state uses the same generation and likelihood protocols, under the instruction *“This is a malignant single cell from a {species}. The gene expression profile is {input}. Which of the following transcriptional meta-programs is most active in this cell: {the* 41 *names}? Answer with only the meta-program name*.*”*. Plain likelihood ranking is strongest here too, and is the InstructCell protocol we report in the main text and in Table 10.

### D.5 Probe on frozen embeddings

This section describes how we probe the single-cell encoder that CellScribe is built on, which gives a discriminative reference on both tasks. The probe keeps that representation fixed and replaces the decoder by a small head, one hidden layer of 256 units with dropout, trained on our training split. For cell typing the head is a classification head over the benchmark label space, trained with cross-entropy, and the prediction is its arg-max. For cell state the head has one output per program and is trained with a multi-label binary cross-entropy against the reference activities binarised at the threshold *τ* =0.5 of the answer template. Its outputs pass through a sigmoid, so each program receives a probability of being active; those above 0.5 form the predicted set, and the full vector of 41 probabilities orders the programs for the ranking metrics. Those probabilities say whether a program is active, not how active it is, which is why the probe has no score MAE either.

## E Cell typing: additional results

This section reports the full typing benchmark of Figure 2a, then the hierarchical metrics for cell typing, and finally the exact numbers for the Tabula Sapiens comparison of Figure 3.

### E.1 The benchmark across training stages

Table 6 reports the benchmark of Figure 2a with the three training stages listed separately rather than as the single CellScribe entry the figure shows.

**Table 6:** Cell typing on the CELLxGENE study-holdout split.

| Model | macro-$F_1$ | weighted-$F_1$ |
| --- | --- | --- |
| scTab (retrained head) | 0.031 | 0.122 |
| CellWhisperer | 0.053 | 0.071 |
| InstructCell | 0.009 | 0.156 |
| Cell2Sentence | 0.083 | 0.356 |
| scFoundation probe | 0.194 | 0.366 |
| CellScribe (SFT) | 0.195 | **0.497** |
| CellScribe (SFT + GRPO) | 0.205 | 0.479 |
| **CellScribe (SFT + GRPO + ICL)** | **0.223** | 0.484 |

### E.2 Hierarchical scoring and label coverage

Exact-match scores charge the same penalty for naming a sibling subtype as for naming an unrelated lineage. Hierarchical *F*_1_ credits the ancestors a prediction shares with the target (Appendix C). Table 7 reports it at four tolerances: 0 edges is exact match, 1 and 2 admit predictions that far up or down the ontology, and *full* keeps every ancestor up to the root, so any shared lineage counts. The ordering of §4 holds at every tolerance and the gap widens rather than closes: CellScribe leads the probe by 2.9 points at exact match and by 3.5 at full lineage, and each of the three stages raises the score at every tolerance. Cell2Sentence, the strongest language baseline at exact match, reaches only 0.194 at full lineage against 0.652 for CellScribe. At tolerance 0 the table reduces to macro-*F*_1_ by construction and reproduces Table 6.

**Table 7:** Hierarchical *F*_1_ on cell typing on CELLxGENE, by ontology tolerance in edges.

| Model | Ontology tolerance (edges) |  |  |  |
| --- | --- | --- | --- | --- |
|  | 0 | 1 | 2 | full |
| scTab (retrained head) | 0.031 | 0.044 | 0.081 | 0.326 |
| CellWhisperer | 0.053 | 0.102 | 0.190 | 0.599 |
| InstructCell | 0.009 | 0.011 | 0.016 | 0.046 |
| Cell2Sentence | 0.083 | 0.100 | 0.125 | 0.194 |
| scFoundation probe | 0.194 | 0.266 | 0.354 | 0.617 |
| CellScribe (SFT) | 0.195 | 0.272 | 0.365 | 0.630 |
| CellScribe (SFT + GRPO) | 0.205 | 0.285 | 0.381 | 0.652 |
| **CellScribe (SFT + GRPO + ICL)** | **0.223** | **0.298** | **0.390** | **0.652** |

### E.3 Cell typing on 3CA

3CA samples come annotated with a cell type as well as a meta-program profile, so the corpus supports the typing task in addition to the state task it is used for in the main text, and we can evaluate cell typing on it too. Its label space is small and cancer-centric compared to CELLxGENE’s: the test split holds 40 types over 661,147 cells, 12 of them tissue-qualified malignant types (`malignant cell (Breast)`, `malignant cell (Brain)`, … ) and 28 microenvironment types such as `T cell`, `fibroblast` and `macrophage`. As explained in Appendix A, all of these types are expressed in the cell ontology used throughout this work. Table 8 evaluates the same three training stages and baselines on that split.

**Table 8:**
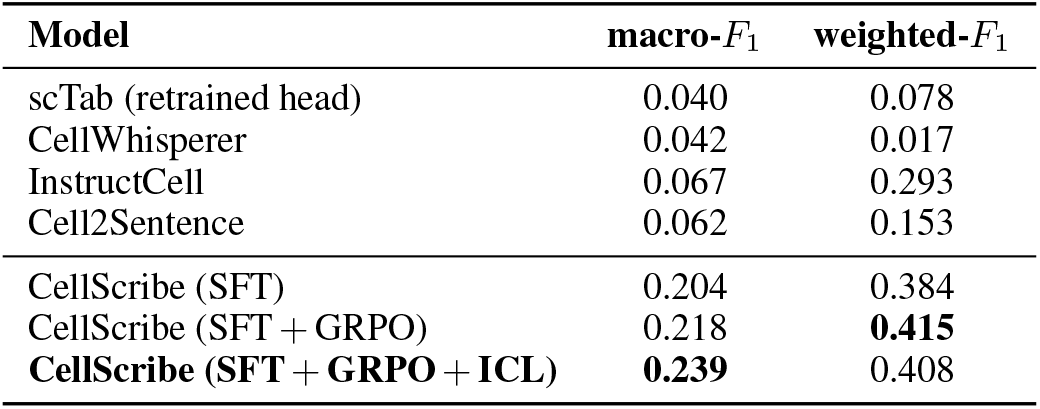
Cell typing on the 3CA test split (40 types, 661,147 cells).

CellScribe leads on both metrics at every stage, more than tripling the best baseline on macro-*F*_1_. The stages behave as they do on CELLxGENE: macro-*F*_1_ rises monotonically across the three, while weighted-*F*_1_ peaks at the alignment stage.

### E.4 Tabula Sapiens

Table 9 reports the benchmark of §4 on Tabula Sapiens, a multi-organ atlas held out from CELLx-GENE training. Its two blocks are two evaluations: the full atlas, and the subset of sixteen types that the main text discusses. On that latter subset each model is run in two inference modes: over its usual vocabulary, as in the first block, and with its answers restricted to the sixteen types, which means constrained decoding over those sixteen names for CellScribe and a candidate list reduced to the same sixteen queries for CellWhisperer.

**Table 9:** Cell typing on Tabula Sapiens. Bold marks the better model within each pair of rows sharing a decision space.

| Evaluation | Model (decision space) | weighted-$F_1$ | accuracy | macro-$F_1$ |
| --- | --- | --- | --- | --- |
| Full atlas (1.14M cells) | CellScribe (open vocabulary) | **0.441** | **0.420** | **0.100** |
|  | CellWhisperer (full candidate list) | 0.072 | 0.057 | 0.014 |
| 16 types (457k cells) | CellScribe (open vocabulary) | **0.713** | **0.635** | **0.522** |
|  | CellWhisperer (full candidate list) | 0.473 | 0.394 | 0.349 |
|  | CellScribe (16 candidates) | **0.780** | **0.775** | **0.624** |
|  | CellWhisperer (16 candidates) | 0.683 | 0.687 | 0.529 |

**Table 10:** Cell state on the 3CA malignant test set. Higher is better except for MAE.

| Model | top-1 | hallmark | set-$F_1$ | NDCG@5 | MAE ↓ |
| --- | --- | --- | --- | --- | --- |
| Cell2Sentence (greedy generation) | 0.022 | 0.069 | 0.027 | 0.025 | — |
| Cell2Sentence (likelihood scoring) | 0.035 | 0.130 | 0.067 | 0.090 | — |
| InstructCell (greedy generation) | 0.020 | 0.094 | 0.023 | 0.022 | — |
| InstructCell (likelihood scoring) | 0.169 | 0.227 | 0.108 | 0.218 | — |
| CellWhisperer (cosine retrieval) | 0.080 | 0.176 | 0.110 | 0.160 | — |
| scFoundation probe | 0.202 | 0.227 | 0.205 | 0.233 | — |
| CellScribe (SFT) | 0.266 | 0.307 | 0.270 | 0.321 | 0.440 |
| CellScribe (SFT + GRPO) | 0.283 | 0.328 | 0.289 | **0.378** | **0.399** |
| **CellScribe (SFT + GRPO + ICL)** | **0.300** | **0.344** | **0.298** | 0.330 | 0.436 |

The sixteen types are: erythrocyte, macrophage, monocyte, classical monocyte, non-classical monocyte, neutrophil, basophil, natural killer cell, CD4-positive alpha-beta T cell, CD8-positive alpha-beta T cell, capillary endothelial cell, pulmonary alveolar type 2 cell, respiratory goblet cell, club cell, basal cell and hepatocyte.

### F Cell-state prediction: additional results

This section reports the cell-state benchmark of Figure 2b with the three training stages listed separately and each baseline shown under every inference protocol of Appendix D.

The discriminative probe, competitive on cell typing (Table 6), falls below every CellScribe stage here, so a discriminative head on the same frozen embedding does not recover a ranked, scored set of co-active programs. Among the baselines, only likelihood scoring over the closed program set is competitive at all: free generation collapses, unsurprisingly, on a vocabulary none of these models was trained on.

## G Design choices

This appendix ablates the three design choices the model builds on: the conditioning mechanism (§G.1), the sampling of cell types during training (§G.2), and the size of the language model (§G.3).

### G.1 Conditioning mechanisms for the scRNA-seq embedding

We compare three ways to pass the scFoundation embedding to the language model: modulating every decoder block (AdaLN-Zero), replacing a `<scRNA-seq>` token with a projected embedding, or expanding the embedding into *K*=16 prompt tokens with a Perceiver [Jaegle et al., 2021] in the resampler form used by Flamingo [Alayrac et al., 2022]. The token mechanism is run in two variants, one where a single linear map produces the injected embedding and one where the projector is parameter-matched to the AdaLN pathway, so that the comparison is not decided by capacity alone (Table 11).

**Table 11:** Conditioning ablations on cell typing and cell state.

| Conditioning | Cell typing |  | Cell state |  |  |  |
| --- | --- | --- | --- | --- | --- | --- |
| | macro-$F_1$ | weighted-$F_1$ | top-1 | hallmark | NDCG@5 | MAE ↓ |
| AdaLN-Zero | **0.195** | **0.497** | **0.266** | **0.307** | **0.321** | **0.440** |
| Perceiver ($K=16$) | 0.174 | 0.367 | 0.118 | 0.136 | 0.129 | 0.709 |
| <scRNA-seq> token, parameter-matched | 0.157 | 0.333 | 0.133 | 0.150 | 0.143 | 0.583 |
| <scRNA-seq> token, simple projection | 0.112 | 0.366 | 0.053 | 0.060 | 0.056 | 1.700 |

### G.2 Class-weighted sampling of cell types

Cell types are distributed with a long tail, so drawing cells in proportion to their frequency would show the model a handful of frequent types almost exclusively. The typing draw is therefore tilted towards rare types by a single exponent, and its effect on cell typing is the claim of this section. Each type *t* carries an inverse-frequency weight

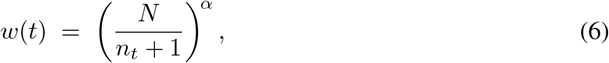

with *N* the number of training cells, *n*_*t*_ the number carrying type *t* and *α* ∈ [0, 1] the exponent we vary. Training reads contiguous chunks of 512 cells; a chunk holding *c*_*t*_ cells of type *t* is drawn with probability proportional to ∑_*t*_ *c*_*t*_ *w*(*t*), and every cell of it is used. A single-type chunk is thus drawn in proportion to 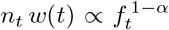 with *f*_*t*_ = *n*_*t*_*/N*, so *α*=0 leaves the natural frequencies, *α*=1 cancels them for a uniform draw over types, and *α*=0.5 gives 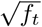. Mixed chunks fall between these, so *α* sets the strength of the correction rather than an exact target distribution. We compare *α*=1 with *α*=0.5 and apply no further cell-level rebalancing.

Table 12 compares the two settings on two runs that share otherwise the same setup and differ only in *α*. They separate exactly as Eq. (6) predicts, and in opposite directions on the two families of typing metric: *α*=1 is 0.9 points ahead on macro-*F*_1_, which weights types equally, while *α*=0.5 is 7.7 and 6.2 points ahead on accuracy and weighted-*F*_1_, which weight cells. We adopt *α*=0.5, the setting favoured by weighted-*F*_1_, the checkpoint-selection metric used throughout, at a cost of 0.9 macro-*F*_1_ points on the tail.

**Table 12:** Class-weighted type sampling under the two values of the exponent *α* of Eq. (6). The *α*=0.5 row is the one the paper builds on.

| Type sampling | Cell typing |  |  | Cell state |  |  |  |
| --- | --- | --- | --- | --- | --- | --- | --- |
| | macro-$F_1$ | acc. | w-$F_1$ | top-1 | hallmark | NDCG@5 | MAE ↓ |
| near-uniform ($\alpha=1$) | **0.205** | 0.401 | 0.435 | 0.224 | 0.253 | 0.252 | 0.458 |
| tempered ($\alpha=0.5$) | 0.195 | **0.478** | **0.497** | 0.266 | 0.307 | 0.321 | 0.440 |

Table 12 also lists cell state for the same two runs, where *α*=0.5 is ahead on every metric. The exponent is a cell-typing lever by design, but it reweights the chunks the model streams, and meta-program annotations are carried only by 3CA cells, so the two runs do not receive the same amount of cell-state supervision. That gain is therefore reported as a property of the two runs rather than attributed to *α* itself; the controlled comparison here is the cell-typing one.

### G.3 Language-model capacity

CellScribe pairs a single-cell foundation encoder with a Qwen2.5-0.5B-Instruct decoder. This section measures the effect of a larger decoder. We vary it in isolation, holding the encoder, the sampler, the budget, the schedule and the AdaLN-Zero conditioning fixed, so the two rows of Table 13 differ in exactly one component.

**Table 13:** Language-model capacity. The two rows differ only in the decoder; the 0.5B row is the one the paper builds on.

| Language model | Cell typing |  |  | Cell state |  |  |
| --- | --- | --- | --- | --- | --- | --- |
| | macro-$F_1$ | acc. | w-$F_1$ | top-1 | hallmark | NDCG@5 |
| Qwen2.5-0.5B | **0.195** | **0.478** | **0.497** | **0.266** | **0.307** | 0.321 |
| Qwen2.5-1.5B | 0.190 | 0.446 | 0.470 | 0.256 | 0.284 | **0.322** |

Moving to Qwen2.5-1.5B-Instruct widens the decoder hidden size from 896 to 1536, which takes the trainable parameters from 79.9M to 269.1M. It costs 3.2 accuracy points and 2.7 weighted-*F*_1_ points on typing and 2.3 hallmark points on state, while macro-*F*_1_ (*−*0.6 points) and NDCG@5 (+0.1 points) are essentially unchanged. Capacity alone therefore buys nothing at this training budget: the larger conditioning stack may need more data or more updates to be trained to the same degree, a direction this work does not pursue. We thus use the 0.5B Qwen decoder throughout this paper.

## H Reward alignment

### H.1 Reward specification

This section details the two rewards used for the reinforcement-learning alignment of CellScribe, one per task.

#### Cell typing

The reward is *R*_type_ = exp(*−d*_*G*_ (*ŷ, y*)), where *d*_*G*_ is the shortest-path distance between the predicted and reference types on the cell ontology graph: an exact match scores 1 and the credit decays with the number of edges separating the two terms. A generation that does not parse as a type receives *−*0.5, and a parsed type absent from the ontology receives *−*1.

#### Cell state

The reward is the mixture of Eq. (2),

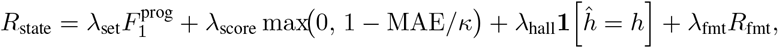

with (*λ*_set_, *λ*_score_, *λ*_hall_, *λ*_fmt_) = (0.35, 0.25, 0.25, 0.15) and *κ* = 3. The answer template of §3 has two parts, the list of active programs and the dominant hallmark, and *R*_fmt_ scores how much of it the generation delivers: 1 when both parts are present and parseable, 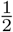 when only one of them is, and 0 when neither is. All four components lie in [0, 1] and their weights sum to one. The coefficients were fixed without a systematic search; measuring the sensitivity of the aligned model to this mixture is left for future work.

### H.2 Dynamic sampling reduces the group skip rate

A group whose *G*=12 completions all receive the same reward has a zero standardised advantage and contributes nothing to the update. Figure 7 measures how often this occurs with and without DAPO-style dynamic sampling [Yu et al., 2025], in which each step oversamples candidate cells threefold, screens each of them with two cheap generations and spends the full twelve-completion rollout only on those whose two screens disagree. The runs differ only in this prescreening, everything else being the same, and it roughly halves the skip rate on both tasks.

**Figure 7:**
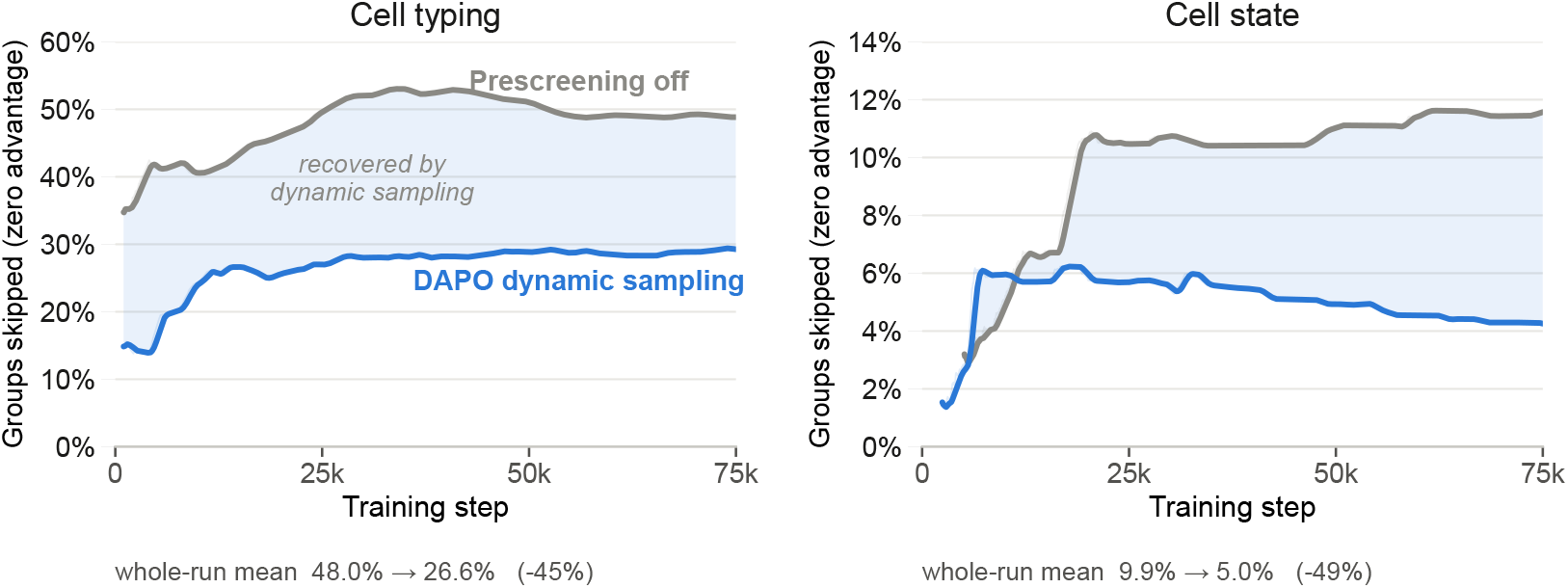
Groups with zero reward variance with prescreening off (grey) or DAPO-style dynamic sampling on (blue). Curves are 100-step rolling means; shading marks the update budget recovered by dynamic sampling.

## I In-context prediction of unseen types

This section reports the exact numbers for the unseen-type experiments of §6, where the target types are excluded from all three training stages and are supplied only as labelled demonstration cells in the prompt. Each of the 4,671 episodes has three reserved candidate types, one labelled support cell per candidate and one query. In Table 14, the rows are the five ways of generating or scoring an answer, and the columns are the models and conditions being compared. We also provide the settings of the in-context post-training.

**Table 14:** Exact-match identification of unseen cell types over the 4,671 episodes. *No surrogate labels* repeats the post-training with real cell type names, and *no ICL post-training* is the reinforcement-learning aligned checkpoint of §5.

| Readout | CellScribe |  | Ablated model |  |
| --- | --- | --- | --- | --- |
|  | correct supports | shuffled supports | no surrogate labels | no ICL post-training |
| Free generation | **0.364** | 0.044 | 0.040 | 0.020 |
| Constrained, full vocabulary | **0.525** | 0.091 | 0.090 | 0.032 |
| Constrained, unseen types (50) | **0.819** | 0.245 | 0.485 | 0.263 |
| Constrained, demonstrated (3) | **0.882** | 0.302 | 0.759 | 0.687 |
| Likelihood ranking (50) | **0.433** | 0.163 | 0.135 | 0.102 |

### In-context learning post-training settings

The stage starts from the aligned checkpoint of §5 and is trained on 6M cells at effective batch 256, of which 80% of the micro-batches are three-way one-shot episodes and 20% are standard single-cell batches mixing cell typing and cell state. We use a cosine learning rate from 6*×*10^*−*5^ to 6*×*10^*−*6^, and the encoder stays frozen. Surrogate labels (§6) are applied to 75% of the episodes.

### Retention of the single-cell tasks

Relative to the aligned checkpoint, typing macro-*F*_1_ rises from 0.205 to 0.223 and hallmark accuracy from 0.328 to 0.344 (Tables 6 and 10); NDCG@5 falls from 0.378 to 0.330. The in-context post-training therefore adds the ability to name unseen types without costing the model its supervised behaviour: on the standard cell typing and cell state benchmarks it leaves CellScribe at least as strong as the checkpoint it starts from on every metric except NDCG@5.

## Notes

### Competing Interest Statement

The authors have declared no competing interest.

## References

Sara Aibar, Carmen Bravo González-Blas, Thomas Moerman, Vân Anh Huynh-Thu, Hana Imrichova, Gert Hulselmans, Florian Rambow, Jean-Christophe Marine, Pierre Geurts, Jan Aerts, et al. SCENIC: single-cell regulatory network inference and clustering. Nature Methods, 14:1083–1086, 2017. doi: 10.1038/nmeth.4463.

Jean-Baptiste Alayrac, Jeff Donahue, Pauline Luc, Antoine Miech, Iain Barr, Yana Hasson, Karel Lenc, Arthur Mensch, Katherine Millican, Malcolm Reynolds, et al. Flamingo: a visual language model for few-shot learning. In Advances in Neural Information Processing Systems, volume 35, pages 23716–23736, 2022.

Jimmy Lei Ba, Jamie Ryan Kiros, and Geoffrey E. Hinton. Layer normalization. arXiv preprint arXiv:1607.06450, 2016.

David A. Barbie, Pablo Tamayo, Jesse S. Boehm, So Young Kim, Susan E. Moody, Ian F. Dunn, Anna C. Schinzel, Peter Sandy, Etienne Meylan, Claudia Scholl, et al. Systematic RNA interference reveals that oncogenic KRAS-driven cancers require TBK1. Nature, 462:108–112, 2009. doi: 10.1038/nature08460.

Dalia Barkley, Reuben Moncada, Maayan Pour, Deborah A. Liberman, Ian Dryg, Gregor Werba, Wei Wang, Maayan Baron, Anjali Rao, Bo Xia, et al. Cancer cell states recur across tumor types and form specific interactions with the tumor microenvironment. Nature Genetics, 54(8):1192–1201, 2022.

Tom B. Brown, Benjamin Mann, Nick Ryder, Melanie Subbiah, Jared Kaplan, Prafulla Dhariwal, Arvind Neelakantan, Pranav Shyam, Girish Sastry, Amanda Askell, et al. Language models are few-shot learners. In Advances in Neural Information Processing Systems, volume 33, pages 1877–1901, 2020.

Haotian Cui, Chloe Wang, Hassaan Maan, Kuan Pang, Fengning Luo, Nan Duan, and Bo Wang. scGPT: toward building a foundation model for single-cell multi-omics using generative AI. Nature Methods, 21(8):1470–1480, 2024.

Sebastiano Cultrera di Montesano, Davide D’Ascenzo, Srivatsan Raghavan, Ava P. Amini, Peter S. Winter, and Lorin Crawford. Improving atlas-scale single-cell annotation models with hierarchical cross-entropy loss. Nature Computational Science, 6:243–249, 2026. doi: 10.1038/s43588-025-00945-z.

CZI Cell Science Program, Shibla Abdulla, Brian Aevermann, Pedro Assis, et al. CZ CELLxGENE Discover: A single-cell data platform for scalable exploration, analysis and modeling of aggregated data. Nucleic Acids Research, 2025.

Alexander D. Diehl, Terrence F. Meehan, Yvonne M. Bradford, Matthew H. Brush, Wasila M. Dahdul, David S. Dougall, Yongqun He, David Osumi-Sutherland, Alan Ruttenberg, Sirarat Sarntivijai, et al. The Cell Ontology 2016: enhanced content, modularization, and ontology interoperability. Journal of Biomedical Semantics, 7:44, 2016. doi: 10.1186/s13326-016-0088-7.

C. Domínguez Conde, C. Xu, L. B. Jarvis, D. B. Rainbow, S. B. Wells, T. Gomes, S. K. Howlett, O. Suchanek, K. Polanski, H. W. King, et al. Cross-tissue immune cell analysis reveals tissue-specific features in humans. Science, 376(6594):eabl5197, 2022.

Yin Fang, Xinle Deng, Kangwei Liu, Ningyu Zhang, Jingyang Qian, Penghui Yang, Xiaohui Fan, and Huajun Chen. A multi-modal AI copilot for single-cell analysis with instruction following. arXiv preprint arXiv:2501.08187, 2025.

Felix Fischer, David S. Fischer, Roman Mukhin, Andrey Isaev, Evan Biederstedt, Alexandra-Chloé Villani and Fabian J. Theis. scTab: Scaling cross-tissue single-cell annotation models. Nature Communications, 15(1):6611, 2024.

Avishai Gavish, Michael Tyler, Alissa C. Greenwald, Rouven Hoefflin, Dor Simkin, Roi Tschernichovsky, Noam Galili Darnell, Einav Somech, Chaya Barbolin, Tomer Antman, et al. Hallmarks of transcriptional intratumour heterogeneity across a thousand tumours. Nature, 618(7965):598–606, 2023.

Robert Geirhos, Jörn-Henrik Jacobsen, Claudio Michaelis, Richard Zemel, Wieland Brendel, Matthias Bethge, and Felix A. Wichmann. Shortcut learning in deep neural networks. Nature Machine Intelligence, 2(11):665–673, 2020. doi: 10.1038/s42256-020-00257-z.

Gemma Team. Gemma 2: Improving open language models at a practical size. arXiv preprint arXiv:2408.00118, 2024.

Minsheng Hao, Jing Gong, Xin Zeng, Chiming Liu, Yucheng Guo, Xingyi Cheng, Taifeng Wang, Jianzhu Ma, Xuegong Zhang, and Le Song. Large-scale foundation model on single-cell transcriptomics. Nature Methods, 21(8):1481–1491, 2024.

Graham Heimberg, Tony Kuo, Daryle J. DePianto, Omar Salem, Tobias Heigl, Nathaniel Diamant, Gabriele Scalia, Tommaso Biancalani, Shannon J. Turley, Jason R. Rock, et al. A cell atlas foundation model for scalable search of similar human cells. Nature, 638:1085–1094, 2025.

Andrew Jaegle, Felix Gimeno, Andy Brock, Oriol Vinyals, Andrew Zisserman, and João Carreira. Perceiver: General perception with iterative attention. In International Conference on Machine Learning, pages 4651–4664. PMLR, 2021.

Kalervo Järvelin and Jaana Kekäläinen. Cumulated gain-based evaluation of IR techniques. ACM Transactions on Information Systems, 20(4):422–446, 2002. doi: 10.1145/582415.582418.

Daniel D. Lee and H. Sebastian Seung. Learning the parts of objects by non-negative matrix factorization. Nature, 401:788–791, 1999. doi: 10.1038/44565.

Jinhyuk Lee, Wonjin Yoon, Sungdong Kim, Donghyeon Kim, Sunkyu Kim, Chan Ho So, and Jaewoo Kang. BioBERT: a pre-trained biomedical language representation model for biomedical text mining. Bioinformatics, 36(4):1234–1240, 2020. doi: 10.1093/bioinformatics/btz682.

Daniel Levine, Syed Asad Rizvi, Sacha Lévy, Nazreen Pallikkavaliyaveetil, David Zhang, Xingyu Chen, Sina Ghadermarzi, Ruiming Wu, Zihe Zheng, Ivan Vrkić, et al. Cell2Sentence: Teaching large language models the language of biology. In Proceedings of the 41st International Conference on Machine Learning (ICML), volume 235 of PMLR, 2024.

Junnan Li, Dongxu Li, Silvio Savarese, and Steven Hoi. BLIP-2: Bootstrapping language-image pre-training with frozen image encoders and large language models. In Proceedings of the 40th International Conference on Machine Learning, pages 19730–19742, 2023.

Ilya Loshchilov and Frank Hutter. Decoupled weight decay regularization. In International Conference on Learning Representations, 2019.

William Peebles and Saining Xie. Scalable diffusion models with transformers. In Proceedings of the IEEE/CVF International Conference on Computer Vision (ICCV), pages 4195–4205, 2023.

Qwen Team. Qwen2.5 technical report. arXiv preprint arXiv:2412.15115, 2024.

Alec Radford, Jong Wook Kim, Chris Hallacy, Aditya Ramesh, Gabriel Goh, Sandhini Agarwal, Girish Sastry, Amanda Askell, Pamela Mishkin, Jack Clark, et al. Learning transferable visual models from natural language supervision. In Proceedings of the 38th International Conference on Machine Learning, pages 8748–8763, 2021.

Colin Raffel, Noam Shazeer, Adam Roberts, Katherine Lee, Sharan Narang, Michael Matena, Yanqi Zhou, Wei Li, and Peter J. Liu. Exploring the limits of transfer learning with a unified text-to-text transformer. Journal of Machine Learning Research, 21(140):1–67, 2020.

Moritz Schaefer, Peter Peneder, Daniel Malzl, Salvo Danilo Lombardo, Mihaela Peycheva, Jake Burton, Anna Hakobyan, Varun Sharma, Thomas Krausgruber, Celine Sin, Jörg Menche, Eleni M. Tomazou, and Christoph Bock. Multimodal learning enables chat-based exploration of single-cell data. Nature Biotechnology, 2025. doi: 10.1038/s41587-025-02857-9.

John Schulman, Filip Wolski, Prafulla Dhariwal, Alec Radford, and Oleg Klimov. Proximal policy optimization algorithms. arXiv preprint arXiv:1707.06347, 2017.

Zhihong Shao, Peiyi Wang, Qihao Zhu, Runxin Xu, Junxiao Song, Xiao Bi, Haowei Zhang, Mingchuan Zhang, Y. K. Li, Y. Wu, and Daya Guo. DeepSeekMath: Pushing the limits of mathematical reasoning in open language models. arXiv preprint arXiv:2402.03300, 2024.

Tim Stuart, Andrew Butler, Paul Hoffman, Christoph Hafemeister, Efthymia Papalexi, William M. Mauck, Yuhan Hao, Marlon Stoeckius, Peter Smibert, and Rahul Satija. Comprehensive integration of single-cell data. Cell, 177(7):1888–1902.e21, 2019. doi: 10.1016/j.cell.2019.05.031.

Fuchou Tang, Catalin Barbacioru, Yangzhou Wang, Ellen Nordman, Clarence Lee, Nanlan Xu, Xiaohui Wang, John Bodeau, Brian B. Tuch, Asim Siddiqui, et al. mRNA-Seq whole-transcriptome analysis of a single cell. Nature Methods, 6:377–382, 2009. doi: 10.1038/nmeth.1315.

The Tabula Sapiens Consortium. The Tabula Sapiens: A multiple-organ, single-cell transcriptomic atlas of humans. Science, 376(6594):eabl4896, 2022.

Christina V. Theodoris, Ling Xiao, Anant Chopra, Mark D. Chaffin, Zeina R. Al Sayed, Matthew C. Hill, Helene Mantineo, Elizabeth M. Brydon, Zexian Zeng, X. Shirley Liu, and Patrick T. Ellinor. Transfer learning enables predictions in network biology. Nature, 618(7965):616–624, 2023.

Itay Tirosh and Mario L. Suvà. Cancer cell states: Lessons from ten years of single-cell RNA-sequencing of human tumors. Cancer Cell, 42(9):1497–1506, 2024.

Sheng Wang, Angela Oliveira Pisco, Aaron McGeever, Maria Brbic, Marinka Zitnik, Spyros Darmanis, Jure Leskovec, Jim Karkanias, and Russ B. Altman. Leveraging the Cell Ontology to classify unseen cell types. Nature Communications, 12(1):5556, 2021.

Jerry Wei, L. Hou, Andrew Lampinen, Xiangning Chen, D. Huang, Yi Tay, Xinyun Chen, Yifeng Lu, Denny Zhou, Tengyu Ma, and Quoc Le. Symbol tuning improves in-context learning in language models. In Proceedings of the 2023 Conference on Empirical Methods in Natural Language Processing, pages 968–979, 2023. doi: 10.18653/v1/2023.emnlp-main.61.

Qiying Yu, Zheng Zhang, Ruofei Zhu, Yufeng Yuan, Xiaochen Zuo, Yu Yue, Weinan Dai, Tiantian Fan, Gaohong Liu, Juncai Liu, et al. DAPO: An open-source LLM reinforcement learning system at scale. In Advances in Neural Information Processing Systems, 2025.

